# Cortical markers of PAS-induced long-term potentiation and depression in the motor system:A TMS-EEG Registered Report

**DOI:** 10.1101/2025.04.03.647045

**Authors:** Eleonora Arrigoni, Nadia Bolognini, Alberto Pisoni, Giacomo Guidali

## Abstract

*Paired associative stimulation* (PAS), a neuromodulation protocol combining transcranial magnetic stimulation (TMS) pulses to the primary motor cortex (M1) with electrical median nerve stimulation, promotes synaptic plasticity (long-term potentiation - LTP, long-term depression - LTD) in the human motor system following Hebbian associative plasticity induction. To date, PAS effects have been mainly investigated at the corticospinal level. In the present Registered Report, we leveraged TMS and electroencephalography (TMS-EEG) co-registration to track the cortical dynamics related to M1-PAS, aiming to characterize the neurophysiological substrates better, grounding the effectiveness of such protocol. In two within-subject sessions, 30 healthy participants underwent the standard M1-PAS protocols inducing LTP (PAS_LTP_) and LTD (PAS_LTD_) while measuring motor-evoked potentials (MEPs) and TMS-evoked potentials (TEPs) from M1 stimulation before, immediately after, and 30 minutes from the end of the PAS, applied both at supra- (i.e., 110%) and sub- (i.e., 90%) resting motor threshold intensities. Besides replicating MEPs enhancement and inhibition after PAS_LTP_ and PAS_LTD_, our results showed that the P30 and N100 M1-TEPs components were significantly modulated immediately following PAS_LTP_ and PAS_LTD_ administration. These effects were detectable only in suprathreshold conditions, suggesting that M1 subthreshold stimulation could not be optimal for tracking cortical effects of PAS. Furthermore, exploratory analyses showed that P60 amplitude at baseline successfully predicted the magnitude of P30 modulations after PAS_LTP_ administration. Our findings provide compelling evidence about the specificity of early TEP components in reflecting changes in M1 reactivity underpinning PAS effects and associative plasticity induction within the motor system. From a broader perspective, our study fosters evidence about using TMS-EEG biomarkers to track complex plastic changes induced in the human brain, exploiting neuromodulatory non-invasive brain stimulation protocols based on associative mechanisms, like PAS.

**Preregistered Stage 1 protocol**: https://osf.io/detjc (date of in-principle acceptance: 15/01/2024)

**Recommended Stage 2 manuscript:** https://rr.peercommunityin.org/articles/rec?id=1031

## 1. INTRODUCTION

*Paired associative stimulation* (PAS) is a class of non-invasive brain stimulation protocols known to induce long-term potentiation (LTP) and long-term depression (LTD) following Hebbian rules of associative plasticity (Hebb, 1949). In PAS protocols, the induction of plasticity is achieved through the repeated pairing of two different stimulations, which activate the same cortical areas or circuits (for a review, see: Suppa et al., 2017).

The standard version of the PAS targets the motor system. It pairs transcranial magnetic stimulation (TMS) pulses over the primary motor cortex (M1) with the electrical stimulation of the contralateral (to TMS) median nerve (M1-PAS) (Stefan et al., 2000). Depending on the inter-stimulus interval (ISI) between these two stimulations, LTP or LTD is induced within the motor system, according to the asymmetric time window of spike-timing-dependent plasticity (STDP) observed in the cellular and animal models (Caporale & Dan, 2008; F. Müller-Dahlhaus et al., 2010). In detail, when the ISI closely resembles the timing in which the afferent sensory signal from the median nerve electrical stimulation reaches M1 (i.e., 25 ms), LTP is induced (PAS_LTP_), with an increase in post-PAS MEPs amplitude (e.g., Conde et al., 2012; Fratello et al., 2006; Nitsche et al., 2007; Stefan et al., 2000; Wolters et al., 2003; Ziemann et al., 2004). Conversely, when the ISI is shorter (i.e., 10 ms) and, thus, the exogenous activation of M1 induced by TMS precedes the endogenous one driven by the electrical stimulation, LTD is induced (PAS_LTD_) (e.g., Batsikadze et al., 2013; Delvendahl et al., 2010; Huber et al., 2008; Stefan et al., 2006; Wolters et al., 2003). The effectiveness of this protocol has been widely replicated in the last two decades (e.g., Kumru et al., 2017; Müller-Dahlhaus et al., 2008; Player et al., 2012; Quartarone et al., 2006; Schabrun et al., 2013) (for reviews, see: Suppa et al., 2017; Wischnewski & Schutter, 2016), and modified versions targeting other cortical areas/networks than M1 and the motor system arose in recent years (e.g., Bevilacqua et al., 2023; Borgomaneri et al., 2023; Casarotto et al., 2023; Di Luzio et al., 2022; Engel et al., 2017; Guidali, Bagattini, et al., 2023; Guidali et al., 2020; Nord et al., 2019; Ranieri et al., 2019; Santarnecchi et al., 2018; Zazio et al., 2019; Zibman et al., 2019) (for reviews, see: Guidali et al., 2021a, 2021b). Proving to be effective tools for inducing LTP/LTD effects, PAS protocols have been extensively used in clinical research to investigate abnormal plasticity in several neuropsychiatric populations (e.g., Brandt et al., 2014; Castel-Lacanal et al., 2009; Crupi et al., 2008; Frantseva et al., 2008; Kuhn et al., 2016; Tolmacheva et al., 2017).

To date, the majority of the studies evaluated the effectiveness of the M1-PAS-induced plasticity within the motor system by focusing, as primary outcomes, on corticospinal excitability (i.e., motor-evoked potentials – MEPs) or behavioral measures (Carson & Kennedy, 2013; Suppa et al., 2017). In the last two decades, concurrent TMS and electroencephalography registration (TMS-EEG) has been extensively used to assess cortical excitability and effective connectivity before and after non-invasive brain stimulation, leveraging the sensitivity of TMS-evoked potentials (TEPs) to track global changes induced by neuromodulation (for reviews, see: Cruciani et al., 2023; Hernandez-Pavon, Veniero, et al., 2023). To the best of our knowledge, up to the present, only two studies (Costanzo et al., 2023; Huber et al., 2008) investigated M1-PAS aftereffects using TMS-EEG.

In a seminal work, Huber and coworkers (2008) measured TMS-evoked activity before and after PAS_LTP_ and PAS_LTD_ to assess modulations of the cortical responses by different ISIs. Results showed that, in single subjects, TMS-evoked cortical responses over sensorimotor cortex changed according to the protocol exploited, representing the first direct evidence that PAS can induce changes in global cortical dynamics. However, in this paper, the authors exploited the global mean field power as the primary variable of interest without analyzing the M1-TEP components profile. Moreover, they qualitatively report differential effects of the two PAS protocols on cortical excitability when applied at different cortical sites, suggesting complex effects of the stimulation protocols on M1 effective connectivity patterns (Huber et al., 2008).

Recently, Costanzo and colleagues (2023) showed that, after the administration of PAS_LTP_, the amplitude of P30 and P60 components of M1-TEPs increased. Different studies highlighted that the P30 reflects local circuits’ excitatory neurotransmission (Bonato et al., 2006; Ferreri et al., 2011; Paus et al., 2001). Along the same line, a P60 modulation was associated with TMS protocols that influence M1 excitability (Esser et al., 2006; Rogasch et al., 2013). No significant correlation was found between increased MEP amplitude and the modulation of single TEP components after the protocol administration. This evidence suggests that peripheral and cortical measures of PAS efficacy frame two different facets of induced plasticity within M1. The study exclusively explored the facilitation effects of PAS (specifically, PAS_LTP_) and analyzed the aftereffects by looking at amplitude modulations of the M1-TEP components only immediately after the protocol’s administration (Costanzo et al., 2023).

Given these premises, in the present study, we aim to deepen the cortical underpinnings of M1-PAS-induced plasticity exploiting TMS-EEG. This investigation is indeed crucial to derive cortical biomarkers of plastic changes in the human brain. To this end, our study aims to better characterize the neurophysiological substrates grounding the effectiveness of non-invasive brain stimulation protocols based on associative mechanisms like PAS ones (e.g., Chung et al., 2015; Ferreri & Rossini, 2013; Kallioniemi & Daskalakis, 2022).

In a within-subjects experiment, healthy participants underwent PAS_LTP_ and PAS_LTD_ protocols (delivered in two different sessions), investigating the spatiotemporal profile of cortical excitability changes (i.e., M1-TEPs) within the motor system before and after the administration of these two M1-PAS protocols. MEPs were recorded as the control variable; namely, we expected that the two protocols would lead to opposite patterns on corticospinal tract excitability, which could be interpreted as LTP- or LTD-like induction within the motor system (Suppa et al., 2017). These patterns served as operative models to discuss the results found on cortico-cortical measures. Indeed, as the *positive control* condition of our study (**H0**), we aim to replicate the corticospinal enhancement and inhibition after PAS_LTP_ and PAS_LTD_, respectively (Wischnewski & Schutter, 2016). Namely, MEPs recorded after PAS_LTP_ are expected to have a greater peak-to-peak amplitude than the ones recorded in baseline, and the opposite pattern should be observed for PAS_LTD_. This analysis would confirm that our two PAS protocols have effectively induced associative plasticity in the expected direction according to previous literature.

Considering PAS effects on early M1-TEP components reflecting local excitability (i.e., P30 and P60 – H1; e.g., Cash et al., 2017; Esser et al., 2006), we expected to replicate, for the PAS_LTP_ protocol, the same pattern of modulation found in the study of Costanzo and coworkers (2023) – i.e., enhancement of P30 and P60 amplitude after excitatory protocol administration. For PAS_LTD_, if LTD induction led to the modulation of the same TEP components, we hypothesized that P30 and P60 would show an amplitude reduction. Notably, these two components are often used as biomarkers of cortical excitability in TMS-EEG studies aimed at assessing the effects of non-invasive neuromodulation techniques inducing LTD/LTP-like phenomena within the motor system (for a review, see: Cruciani et al., 2023).

In detail, P30 is thought to reflect fast excitatory mechanisms within M1 local circuitry (Mäki & Ilmoniemi, 2010; Rogasch et al., 2013). Hence, P30 was reported to be positively correlated with MEP amplitude (Ferreri et al., 2011; Mäki & Ilmoniemi, 2010). Corroborating this hypothesis, intermittent (iTBS) and continuous (cTBS) theta-burst TMS – used to transiently increase and suppress motor cortex excitability, respectively – influence P30 amplitude in the same direction of MEP modulations. For instance, inhibition of P30 was found following cTBS (Vernet et al., 2013), and Gedankien and colleagues (2017) showed that iTBS-induced changes in N15-P30 TEP complex and MEP amplitude were significantly correlated (Gedankien et al., 2017). On the other hand, P60 has been associated with the activity of recurrent cortico-cortical and cortico-subcortical circuits reflecting glutamatergic signal propagation mediated by AMPA receptor activation (Belardinelli et al., 2021). Previous TMS-EEG evidence showed that the P60 component can be modulated by drugs influencing gamma-aminobutyric acid (GABA) neurotransmission (Gordon et al., 2022), suggesting that the P60 amplitude likely reflects the excitation/inhibition balance of the stimulated region. In fact, different TMS and transcranial direct current stimulation interventions significantly modulated the amplitude of the TMS-evoked P60 after their application (Chung et al., 2019; Mosayebi-Samani et al., 2023).

Considering later M1-TEP components (**H2**), it is well known that the N100 is a marker of inhibitory processing mediated by GABA receptors, and different studies related the modulation of this component to the induction of inhibitory-like phenomena or plastic effects (Bonnard et al., 2009; Casula et al., 2014; Premoli et al., 2018; Premoli, Rivolta, et al., 2014; Rogasch et al., 2013). Similarly, we expected that the N100 would be influenced by PAS_LTD_ administration. Hence, considering the inhibitory nature of this component, we hypothesized that PAS_LTD_ administration would lead to a greater (negative) amplitude of this component. Noteworthy is that Costanzo et al. (2023) found no significant modulation of the N100 after PAS_LTP_. So, given the controversial literature on N100 modulations after the administration of excitatory TMS protocols (e.g., Bai et al., 2021; Chung et al., 2019; Desforges et al., 2022; Goldsworthy et al., 2020), no analysis on PAS_LTP_- N100 effects were registered.

Then, we have deepened the duration of PAS aftereffects on cortical excitability (**H3**). Namely, whether PAS modulations recorded at a cortical level exhibited the same temporal evolution as the effects typically observed on MEPs. To this aim, MEPs and TEPs were assessed 30 minutes after the PAS administration. Previous studies showed that PAS aftereffects are detectable in a time window of about double the time of protocol duration (Suppa et al., 2017; Wischnewski & Schutter, 2016; Wolters et al., 2003). Hence, based on previous evidence and considering that our PAS protocols lasted 15 minutes (see **2.3**), we hypothesized that induced plasticity patterns fade away about 30 minutes after the end of the protocol, likely for both PAS_LTP_ and PAS_LTD_.

If this is true, we expected a significant difference to emerge when comparing TMS-evoked activity (i.e., P30, P60, N100, and MEP amplitude) after the intervention with the one recorded after 30 minutes.

Finally, different studies argued that the interpretation of the functional meaning of P60 might be possibly hampered by confounding factors related to the elaboration of afferent proprioceptive signals related to MEPs (i.e., P60; e.g., Fecchio et al., 2017; Komssi et al., 2004) with respect to early components (i.e., P30; e.g., Gordon et al., 2018; Petrichella et al., 2017). This aspect complicates the interpretation of P60, making it difficult to disentangle the contribution of peripheral processing to the amplitude increases of this cortical component found after PAS. In detail, as previously noted for PAS_LTP_ (Costanzo et al., 2023), we hypothesized that, in such a protocol, the change in P60 magnitude could be overestimated due to the involvement of MEP reafference (**H4**). Hence, to rule out this hypothesis and provide more detailed information for the overall interpretation of the results, before and after PAS administration, M1-TEPs were recorded at a subthreshold intensity (i.e., 90% of participant’s resting motor threshold – rMT), besides being recorded at a standard suprathreshold intensity (i.e., 110% rMT). If the reafferent signals have a major impact on P60 amplitude modulation, we expected that, compared to P30 (which is too early and allegedly unaffected by MEP reafference), P60 would display a greater change in amplitude in the suprathreshold condition after PAS_LTP_ due to the MEP presence. Noteworthy, previous literature showed that TEPs could be successfully recorded at subthreshold intensities, displaying the same typical components as suprathreshold TEPs (Komssi et al., 2004; Lioumis et al., 2009). Given the rationale behind this fourth hypothesis, we would test it only if a significant modulation of P60 is found in **H1**.

Overall, our study aimed to explore possible cortical markers of Hebbian associative LTP- and LTD-like plasticity in the motor system exploiting the PAS protocol. This investigation took advantage of concurrent TMS-EEG registration, deepening the spatiotemporal patterns of M1-TEPs after the administration of excitatory and inhibitory M1-PAS protocols (see **Table 1** for all our *a priori* hypotheses and related planned analysis).

**Table 1.**
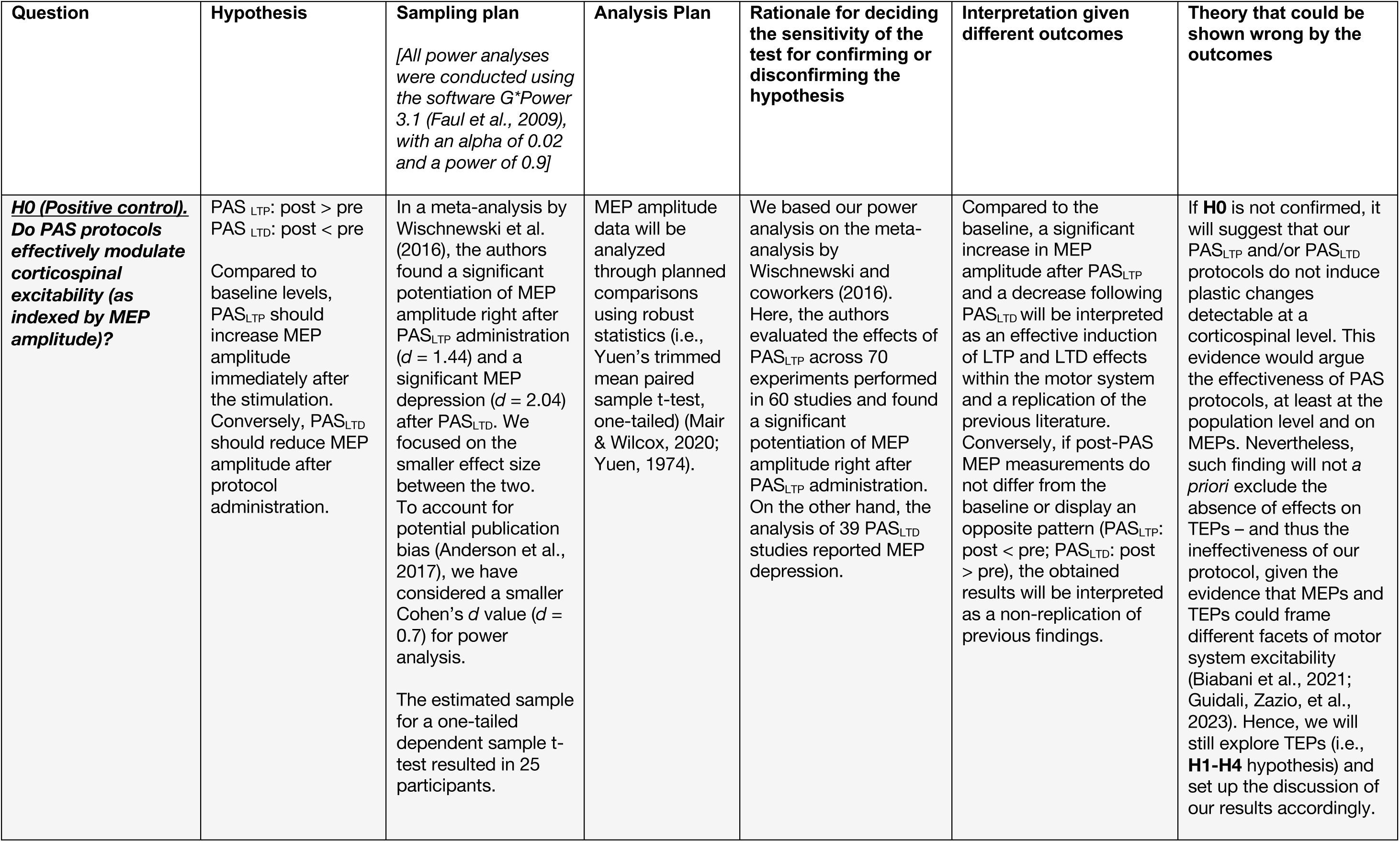
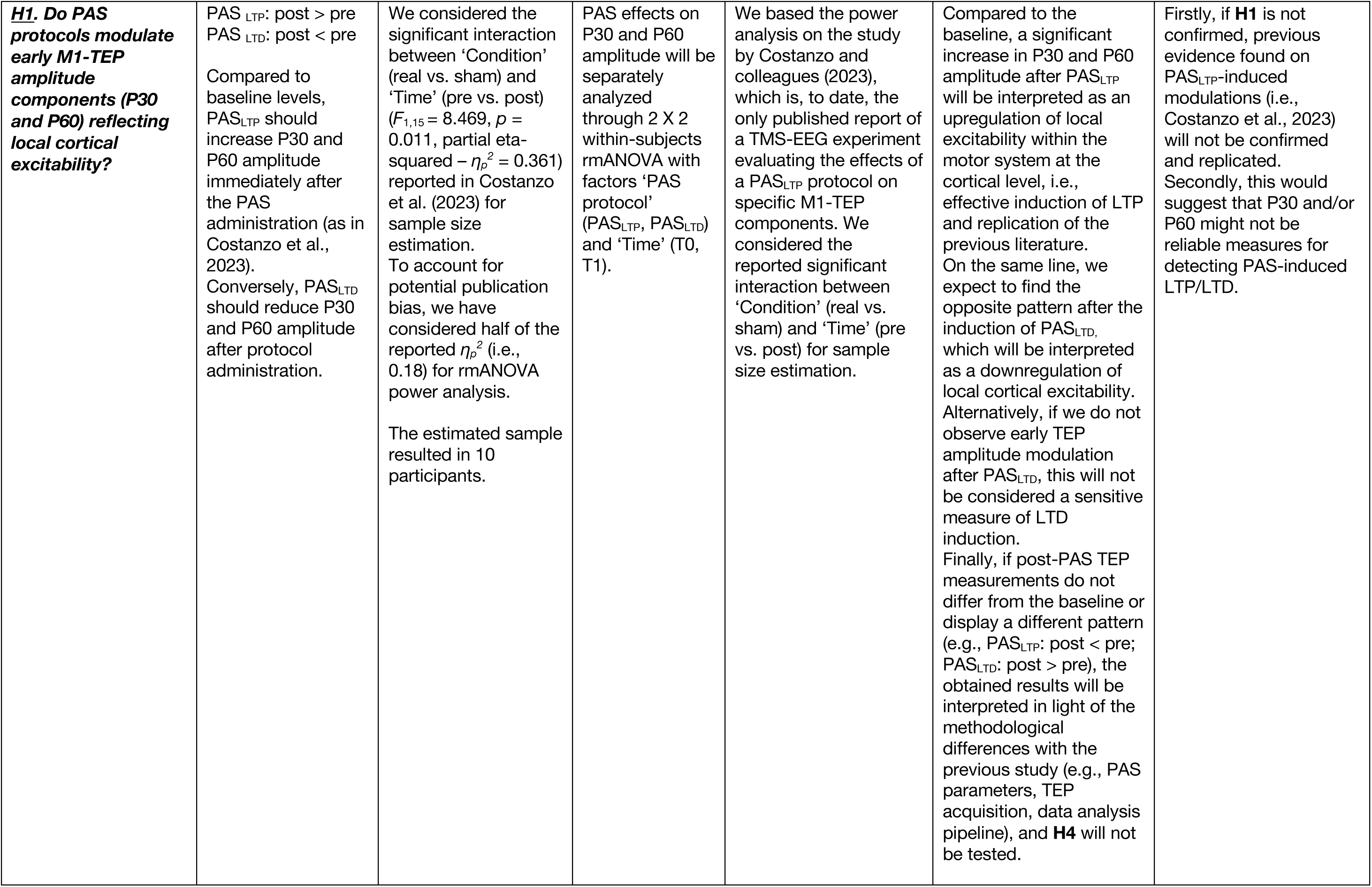
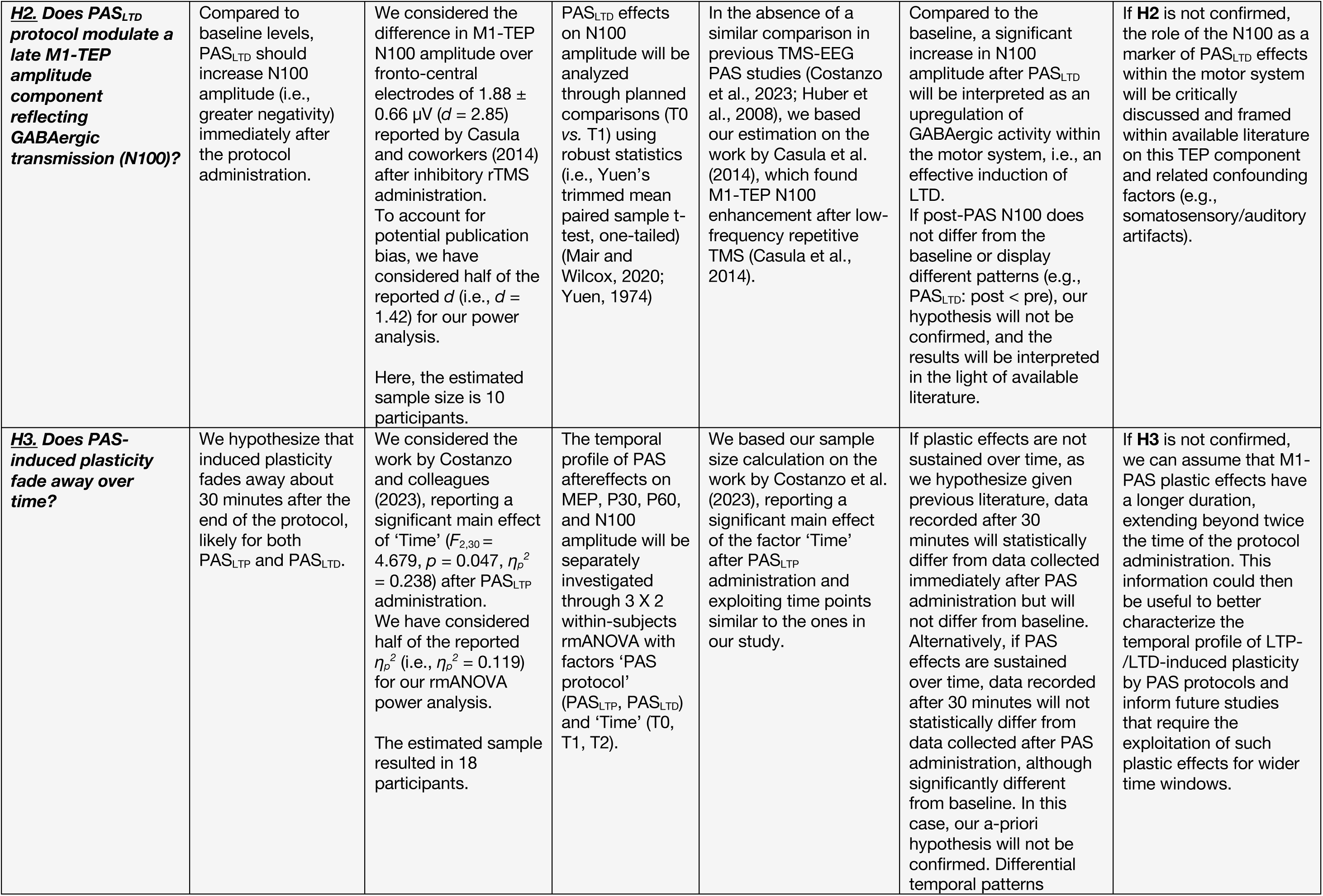
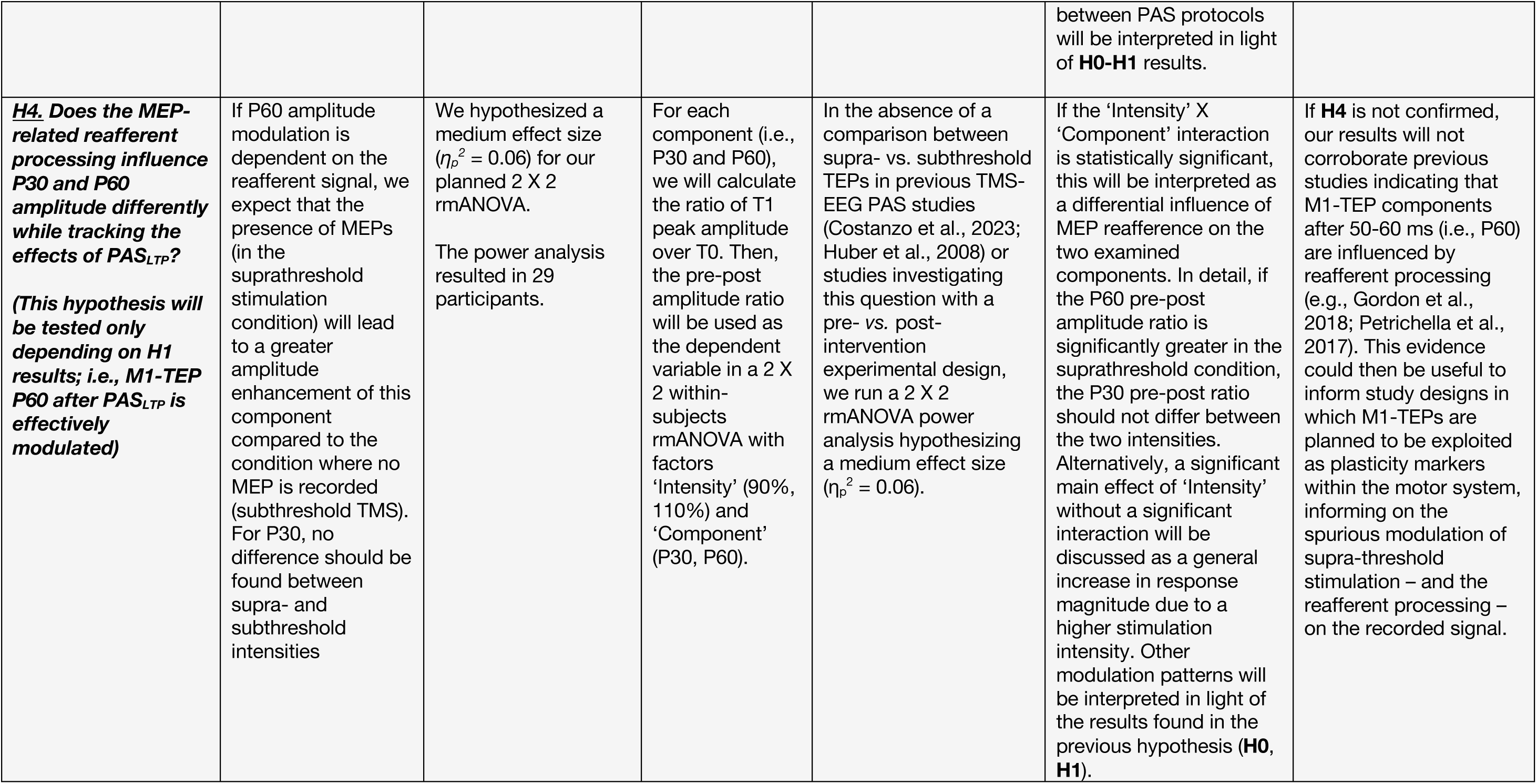
Study design.

| Question | Hypothesis | Sampling plan<br>*[All power analyses were conducted using the software G\*Power 3.1 (Faul et al., 2009), with an alpha of 0.02 and a power of 0.9]* | Analysis Plan | Rationale for deciding the sensitivity of the test for confirming or disconfirming the hypothesis | Interpretation given different outcomes | Theory that could be shown wrong by the outcomes |
| --- | --- | --- | --- | --- | --- | --- |
| **<u>H0 (Positive control).</u>**<br>**Do PAS protocols effectively modulate corticospinal excitability (as indexed by MEP amplitude)?** | PAS<sub>LTP</sub>: post > pre<br>PAS<sub>LTD</sub>: post < pre<br>Compared to baseline levels, PAS<sub>LTP</sub> should increase MEP amplitude immediately after the stimulation. Conversely, PAS<sub>LTD</sub> should reduce MEP amplitude after protocol administration. | In a meta-analysis by Wischniewski et al. (2016), the authors found a significant potentiation of MEP amplitude right after PAS<sub>LTP</sub> administration ($d = 1.44$) and a significant MEP depression ($d = 2.04$) after PAS<sub>LTD</sub>. We focused on the smaller effect size between the two. To account for potential publication bias (Anderson et al., 2017), we have considered a smaller Cohen's $d$ value ($d = 0.7$) for power analysis.<br>The estimated sample for a one-tailed dependent sample t-test resulted in 25 participants. | MEP amplitude data will be analyzed through planned comparisons using robust statistics (i.e., Yuen's trimmed mean paired sample t-test, one-tailed) (Mair & Wilcox, 2020; Yuen, 1974). | We based our power analysis on the meta-analysis by Wischniewski and coworkers (2016). Here, the authors evaluated the effects of PAS<sub>LTP</sub> across 70 experiments performed in 60 studies and found a significant potentiation of MEP amplitude right after PAS<sub>LTP</sub> administration. On the other hand, the analysis of 39 PAS<sub>LTD</sub> studies reported MEP depression. | Compared to the baseline, a significant increase in MEP amplitude after PAS<sub>LTP</sub> and a decrease following PAS<sub>LTD</sub> will be interpreted as an effective induction of LTP and LTD effects within the motor system and a replication of the previous literature. Conversely, if post-PAS MEP measurements do not differ from the baseline or display an opposite pattern (PAS<sub>LTP</sub>: post < pre; PAS<sub>LTD</sub>: post > pre), the obtained results will be interpreted as a non-replication of previous findings. | If **H0** is not confirmed, it will suggest that our PAS<sub>LTP</sub> and/or PAS<sub>LTD</sub> protocols do not induce plastic changes detectable at a corticospinal level. This evidence would argue the effectiveness of PAS protocols, at least at the population level and on MEPs. Nevertheless, such finding will not *a priori* exclude the absence of effects on TEPs – and thus the ineffectiveness of our protocol, given the evidence that MEPs and TEPs could frame different facets of motor system excitability (Biabani et al., 2021; Guidali, Zazio, et al., 2023). Hence, we will still explore TEPs (i.e., **H1-H4** hypothesis) and set up the discussion of our results accordingly. |
| **<u>H1. Do PAS protocols modulate early M1-TEP amplitude components (P30 and P60) reflecting local cortical excitability?</u>** | PAS<sub>LTP</sub>: post > pre<br>PAS<sub>LTD</sub>: post < pre<br>Compared to baseline levels, PAS<sub>LTP</sub> should increase P30 and P60 amplitude immediately after the PAS administration (as in Costanzo et al., 2023). Conversely, PAS<sub>LTD</sub> should reduce P30 and P60 amplitude after protocol administration. | We considered the significant interaction between ‘Condition’ (real vs. sham) and ‘Time’ (pre vs. post) ($F_{1,15} = 8.469$, $p = 0.011$, partial eta-squared – $\eta_p^2 = 0.361$) reported in Costanzo et al. (2023) for sample size estimation. To account for potential publication bias, we have considered half of the reported $\eta_p^2$ (i.e., 0.18) for rmANOVA power analysis.<br>The estimated sample resulted in 10 participants. | PAS effects on P30 and P60 amplitude will be separately analyzed through 2 X 2 within-subjects rmANOVA with factors ‘PAS protocol’ (PAS<sub>LTP</sub>, PAS<sub>LTD</sub>) and ‘Time’ (T0, T1). | We based the power analysis on the study by Costanzo and colleagues (2023), which is, to date, the only published report of a TMS-EEG experiment evaluating the effects of a PAS<sub>LTP</sub> protocol on specific M1-TEP components. We considered the reported significant interaction between ‘Condition’ (real vs. sham) and ‘Time’ (pre vs. post) for sample size estimation. | Compared to the baseline, a significant increase in P30 and P60 amplitude after PAS<sub>LTP</sub> will be interpreted as an upregulation of local excitability within the motor system at the cortical level, i.e., effective induction of LTP and replication of the previous literature. On the same line, we expect to find the opposite pattern after the induction of PAS<sub>LTD</sub>, which will be interpreted as a downregulation of local cortical excitability. Alternatively, if we do not observe early TEP amplitude modulation after PAS<sub>LTD</sub>, this will not be considered a sensitive measure of LTD induction. Finally, if post-PAS TEP measurements do not differ from the baseline or display a different pattern (e.g., PAS<sub>LTP</sub>: post < pre; PAS<sub>LTD</sub>: post > pre), the obtained results will be interpreted in light of the methodological differences with the previous study (e.g., PAS parameters, TEP acquisition, data analysis pipeline), and **H4** will not be tested. | Firstly, if **H1** is not confirmed, previous evidence found on PAS<sub>LTP</sub>-induced modulations (i.e., Costanzo et al., 2023) will not be confirmed and replicated. Secondly, this would suggest that P30 and/or P60 might not be reliable measures for detecting PAS-induced LTP/LTD. |
| ***H2. Does PAS<sub>LTD</sub> protocol modulate a late M1-TEP amplitude component reflecting GABAergic transmission (N100)?*** | Compared to baseline levels, PAS<sub>LTD</sub> should increase N100 amplitude (i.e., greater negativity) immediately after the protocol administration. | We considered the difference in M1-TEP N100 amplitude over fronto-central electrodes of $1.88 \pm 0.66 \mu V$ ($d = 2.85$) reported by Casula and coworkers (2014) after inhibitory rTMS administration. To account for potential publication bias, we have considered half of the reported $d$ (i.e., $d = 1.42$) for our power analysis.<br>Here, the estimated sample size is 10 participants. | PAS<sub>LTD</sub> effects on N100 amplitude will be analyzed through planned comparisons (T0 vs. T1) using robust statistics (i.e., Yuen's trimmed mean paired sample t-test, one-tailed) (Mair and Wilcox, 2020; Yuen, 1974) | In the absence of a similar comparison in previous TMS-EEG PAS studies (Costanzo et al., 2023; Huber et al., 2008), we based our estimation on the work by Casula et al. (2014), which found M1-TEP N100 enhancement after low-frequency repetitive TMS (Casula et al., 2014). | Compared to the baseline, a significant increase in N100 amplitude after PAS<sub>LTD</sub> will be interpreted as an upregulation of GABAergic activity within the motor system, i.e., an effective induction of LTD. If post-PAS N100 does not differ from the baseline or display different patterns (e.g., PAS<sub>LTD</sub>: post < pre), our hypothesis will not be confirmed, and the results will be interpreted in the light of available literature. | If **H2** is not confirmed, the role of the N100 as a marker of PAS<sub>LTD</sub> effects within the motor system will be critically discussed and framed within available literature on this TEP component and related confounding factors (e.g., somatosensory/auditory artifacts). |
| ***H3. Does PAS-induced plasticity fade away over time?*** | We hypothesize that induced plasticity fades away about 30 minutes after the end of the protocol, likely for both PAS<sub>LTP</sub> and PAS<sub>LTD</sub>. | We considered the work by Costanzo and colleagues (2023), reporting a significant main effect of 'Time' ($F_{2,30} = 4.679$, $p = 0.047$, $\eta_p^2 = 0.238$) after PAS<sub>LTP</sub> administration. We have considered half of the reported $\eta_p^2$ (i.e., $\eta_p^2 = 0.119$) for our rmANOVA power analysis.<br>The estimated sample resulted in 18 participants. | The temporal profile of PAS aftereffects on MEP, P30, P60, and N100 amplitude will be separately investigated through 3 X 2 within-subjects rmANOVA with factors 'PAS protocol' (PAS<sub>LTP</sub>, PAS<sub>LTD</sub>) and 'Time' (T0, T1, T2). | We based our sample size calculation on the work by Costanzo et al. (2023), reporting a significant main effect of the factor 'Time' after PAS<sub>LTP</sub> administration and exploiting time points similar to the ones in our study. | If plastic effects are not sustained over time, as we hypothesize given previous literature, data recorded after 30 minutes will statistically differ from data collected immediately after PAS administration but will not differ from baseline. Alternatively, if PAS effects are sustained over time, data recorded after 30 minutes will not statistically differ from data collected after PAS administration, although significantly different from baseline. In this case, our a-priori hypothesis will not be confirmed. Differential temporal patterns<br>between PAS protocols will be interpreted in light of **H0-H1** results. | If **H3** is not confirmed, we can assume that M1-PAS plastic effects have a longer duration, extending beyond twice the time of the protocol administration. This information could then be useful to better characterize the temporal profile of LTP-/LTD-induced plasticity by PAS protocols and inform future studies that require the exploitation of such plastic effects for wider time windows. |
| ***H4. Does the MEP-related reafferent processing influence P30 and P60 amplitude differently while tracking the effects of PAS<sub>LTP</sub>?***<br>***(This hypothesis will be tested only depending on H1 results; i.e., M1-TEP P60 after PAS<sub>LTP</sub> is effectively modulated)*** | If P60 amplitude modulation is dependent on the reafferent signal, we expect that the presence of MEPs (in the suprathreshold stimulation condition) will lead to a greater amplitude enhancement of this component compared to the condition where no MEP is recorded (subthreshold TMS). For P30, no difference should be found between supra- and subthreshold intensities | We hypothesized a medium effect size ($\eta_p^2 = 0.06$) for our planned 2 X 2 rmANOVA.<br>The power analysis resulted in 29 participants. | For each component (i.e., P30 and P60), we will calculate the ratio of T1 peak amplitude over T0. Then, the pre-post amplitude ratio will be used as the dependent variable in a 2 X 2 within-subjects rmANOVA with factors 'Intensity' (90%, 110%) and 'Component' (P30, P60). | In the absence of a comparison between supra- vs. subthreshold TEPs in previous TMS-EEG PAS studies (Costanzo et al., 2023; Huber et al., 2008) or studies investigating this question with a pre- vs. post-intervention experimental design, we run a 2 X 2 rmANOVA power analysis hypothesizing a medium effect size ($\eta_p^2 = 0.06$). | If the 'Intensity' X 'Component' interaction is statistically significant, this will be interpreted as a differential influence of MEP reafference on the two examined components. In detail, if the P60 pre-post amplitude ratio is significantly greater in the suprathreshold condition, the P30 pre-post ratio should not differ between the two intensities. Alternatively, a significant main effect of 'Intensity' without a significant interaction will be discussed as a general increase in response magnitude due to a higher stimulation intensity. Other modulation patterns will be interpreted in light of the results found in the previous hypothesis (**H0**, **H1**). | If **H4** is not confirmed, our results will not corroborate previous studies indicating that M1-TEP components after 50-60 ms (i.e., P60) are influenced by reafferent processing (e.g., Gordon et al., 2018; Petrichella et al., 2017). This evidence could then be useful to inform study designs in which M1-TEPs are planned to be exploited as plasticity markers within the motor system, informing on the spurious modulation of supra-threshold stimulation – and the reafferent processing – on the recorded signal. |

## 2. MATERIALS AND METHODS

### 2.1. Participants

Healthy participants (age range: 18-40 years) were recruited for the present study. All participants were right-handed, as assessed with the Edinburgh handedness questionnaire (Oldfield, 1971), with no contraindications to TMS administration following TMS safety guidelines (Rossi et al., 2021) and no history of neurological, psychiatric, or other relevant medical conditions. Participants taking medications known to affect PAS effects (i.e., corticosteroids, anxiolytics, centrally acting ion channel blockers, or antihistamines) were a-priori excluded from the study unless, at the time of the first session of the experiment, they had not taken such medications for at least one month before the assessment (Suppa et al., 2017). Each participant completed a safety screening questionnaire to exclude the presence of contraindications to TMS (Rossi et al., 2021) and gave informed written consent before participating in the study. The study was performed in the TMS-EEG laboratory of the University of Milano-Bicocca following the Declaration of Helsinki and has received approval from the local Ethics Committee (protocol number 797-23). All participants belong to the same experimental group and underwent the same procedures. Participants were naïve to the testing procedures and were debriefed immediately after the end of the last session. Detailed information on the final sample size is reported in section **3.1**.

### 2.2. Sample size estimation

Here, we provide the rationale for the sample size estimation of each experimental hypothesis (**Table 1**). All the analyses were conducted using the software G*Power 3.1 (Faul et al., 2009), with an alpha of 0.02 and a power of 0.9. Of all of them, we ultimately considered the largest sample size for the present study.

#### a) H0 (positive control): Effects of PAS protocols on MEP amplitude

For the *positive control* of our study, we based our sample size estimation on a meta-analysis by Wischnewski and colleagues (2016). Here, the authors evaluated the effects of PAS_LTP_ across 70 experiments performed in 60 studies and found a significant potentiation of corticospinal output (as indexed by MEPs amplitude) right after protocol administration (Cohen’s *d* = 1.44). On the other hand, the analysis of 39 PAS_LTD_ studies demonstrated a consistent depression of cortical excitability levels compared to baseline immediately after this M1-PAS version (*d* = 2.04). We used information from this meta-analysis to retrieve Cohen’s *d* values for the planned t-tests and focused on the smaller effect size between the two (i.e., *d* = 1.44). To account for potential publication bias (Anderson et al., 2017), we have considered half of the reported Cohen’s *d* value (*d* = 0.7) for power analysis. Hence, the estimated sample for a one-tailed dependent sample t-test resulted in 25 participants.

#### b) H1: Effects of PAS protocols on early positive TEP components (P30 and P60)

Concerning the effects of PAS on early TEPs (i.e., P30 and P60), we considered the study by Costanzo et al. (2023), which is, to date, the only published report of a TMS-EEG experiment evaluating the effects of a PAS_LTP_ protocol on these specific M1-TEP components. From this article, we considered the reported significant interaction between ‘Condition’ and ‘Time’ factors (*F*_3,45_ = 8.469, *p* = 0.011, partial eta-squared – *η_p_^2^* = 0.361) for our sample size estimation (Costanzo et al., 2023). As for the previous estimation, we have considered half of the reported *η_p_^2^* (i.e., *η_p_^2^* = 0.18) for a 2 X 2 rmANOVA power analysis to account for potential publication bias. The estimated sample resulted in 10 participants.

#### c) H2: Effects of PAS_LTD_ on the N100

Based on previous literature about LTD and M1-TEPs (Casula et al., 2014), and in the absence of a similar comparison in previous TMS-EEG PAS studies (Costanzo et al., 2023; Huber et al., 2008), we based our estimation on the work by Casula et al. (2014) which found M1-TEP N100 enhancement after low-frequency (i.e., inhibitory) repetitive TMS. The authors reported a difference in N100 amplitude over fronto-central electrodes of 1.88 ± 0.66 μV corresponding to a Cohen’s *d* of 2.85 (Casula et al., 2014). As for the previous estimations, we have considered half of the reported *d* (i.e., *d* = 1.42) for our power analysis to account for potential publication bias. Here, the estimated sample size for a one-tailed dependent sample t-test is 10 participants.

#### d) H3: Temporal evolution of induced plasticity

Here, we will evaluate the temporal evolution of the two PAS protocols. Sample size estimation is based on the work by Costanzo and colleagues (2023), reporting a significant main effect of ‘Time’ (*F*_2,30_ = 4.679, *p* = 0.047, *η_p_^2^* = 0.238) after PAS_LTP_ administration and exploiting timepoints similar to the ones of our study. As for the previous estimations, we have considered half of the reported *η_p_^2^* (i.e., *η_p_^2^* = 0.119) for our rmANOVA power analysis to account for potential publication bias. The estimated sample was found to be 18 participants.

#### e) H4: Effects of TMS pulse intensity on the modulation of P30 and P60 after PAS_LTP_

Finally, our study will examine P30 and P60 modulations elicited by supra- and subthreshold TMS pulses after PAS_LTP_. Considering only the excitatory version of the M1-PAS, in the absence of comparison between supra- *vs.* subthreshold TEPs in previous TMS-EEG PAS studies (Costanzo et al., 2023; Huber et al., 2008), as well as in previous TMS-EEG literature testing the effects of stimulation intensity in a pre- *versus* post-intervention experimental design as ours, we run a 2 X 2 rmANOVA power analysis hypothesizing a medium effect size (*η_p_^2^ =* 0.06) (Fritz et al., 2012). Notably, given the effect sizes found in previous literature that has explored M1-TEP modulations by applying TMS below or above the individual rMT (Lioumis et al., 2009), as well as in trials with or without MEPs (Petrichella et al., 2017), this value is configured as sufficient to detect statistically significant effects of interest. Here, the estimated sample size is 29 participants.

2.3. Taken together all the sample size estimations for our hypotheses, 30 participants would be recruited for the study to allow proper counterbalancing of the experimental conditions. Additional participants would be recruited if needed to compensate for the possibility of dropouts or outliers (see **2.3**) until the required number of 30 complete datasets was reached.

### 2.4. Exclusion criteria

Participants were excluded from the study if one of the following criteria was met:

a. Participants failed the initial screening – i.e., they resulted left-handed on the Edinburgh questionnaire (score below 0), presented contraindications to TMS according to Rossi et al.’s (2021) safety guidelines, or made chronic/acute use of PAS-influencing medications as reported in **2.1**.
b. Participants did not complete all the experimental procedures or both sessions.
c. TMS intensity exceeded 80% of the maximum stimulator output in at least one session.
d. MEP amplitude, TEP P30, P60, and N100 amplitude exceeded 3 SD from the group mean in at least one recording block.
e. More than 10% of the EEG channels were marked as bad (i.e., broken, excessive noise) by visual inspection of the trials during TMS-EEG preprocessing in at least one of the recording blocks.
f. Less than 20 TMS-EMG trials or 80 TMS-EEG trials survived after trial rejection during preprocessing in at least one of the recording blocks.
g. TMS-EEG cleaned data had a low signal-to-noise ratio – SNR (< 1.5) defined as the ratio of mean absolute amplitude of EEG during the 300 ms post-TMS period over the range of the baseline amplitude.

### 2.5. Experimental procedure

The study consisted of a within-subjects design in two sessions separated by a washout period of at least one week to avoid PAS carry-over effects (Suppa et al., 2017). The two sessions were carried out at the same moment (i.e., in the morning or the afternoon). Participants sat comfortably in a semi-reclined armchair in front of a 20” computer screen at a distance of 100 cm, with their arms relaxed on the armrests. All the experimental procedures were the same between the two sessions, except for the PAS protocol administered (i.e., PAS_LTP_ or PAS_LTD_). As in Huber et al. (2008), we decided not to introduce a sham condition because previous PAS literature already provides substantial evidence about the difference in the effective outcomes of the two exploited protocols, at least considering MEP modulations (Wischnewski & Schutter, 2016).

Experimental procedures are summarized in **Figure 1**. Before each experimental session, the motor hotspot of the right *abductor pollicis brevis* (APB) muscle (stimulation target) was localized through neuronavigation procedures, and rMT was determined (see **2.5**).

**Figure 1.**
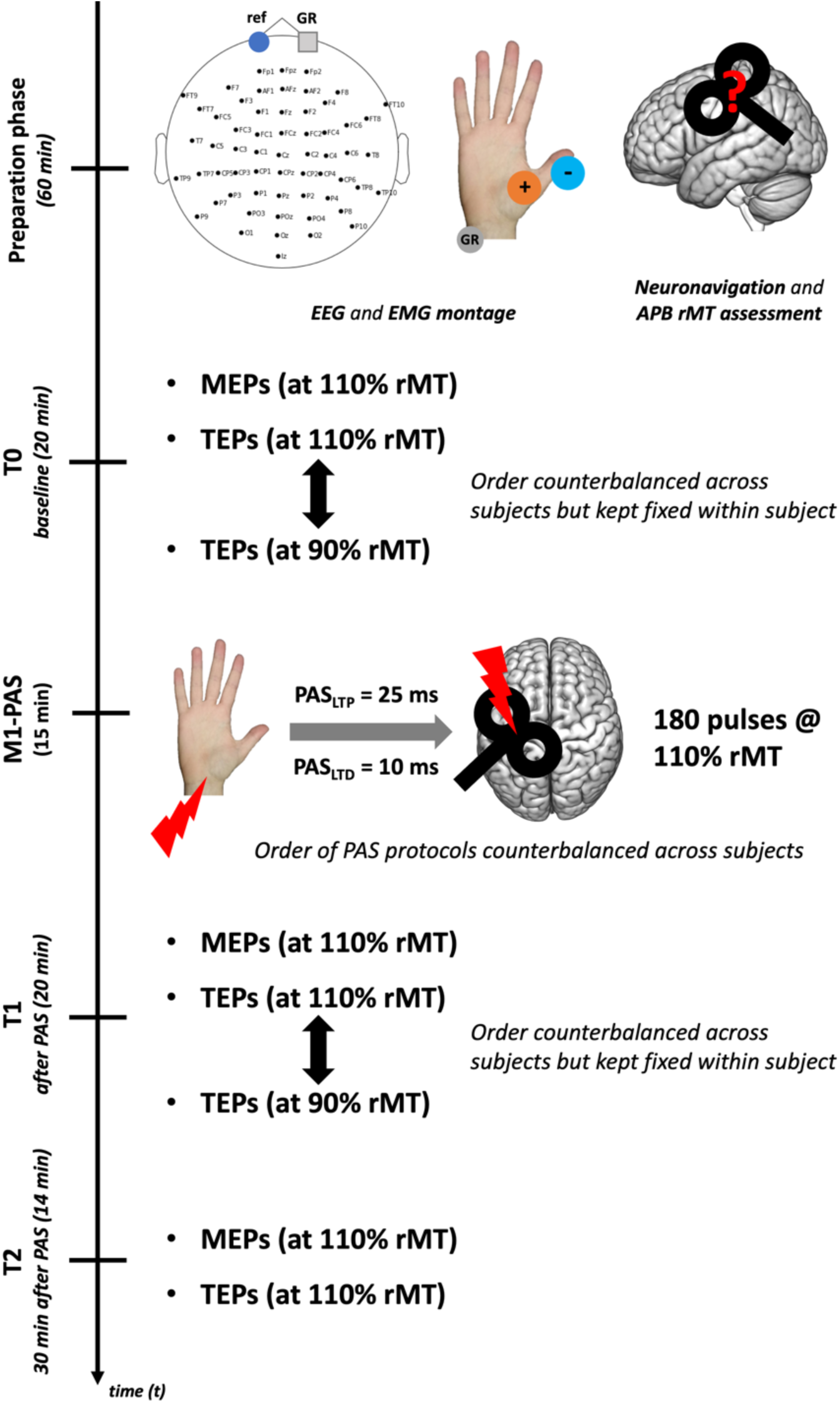
Experimental procedure. After EEG and EMG montage, neuronavigation procedures and APB motor hotspot assessment were carried out. Then, baseline (i.e., T0) MEPs and TEPs at supra- (110% rMT) and sub- threshold (90%) intensity were recorded. After this initial assessment, M1-PAS was administered, and MEPs/TEPs were re-assessed immediately after (i.e., T1) and 30 minutes after (i.e., T2) protocol’s administration. At T2, only TEPs at supra-threshold intensity were recorded. TMS was administered over the left M1, keeping all stimulation parameters constant throughout all our experimental blocks.

PAS protocols were performed by pairing electrical median nerve stimulation with TMS over the left M1, as in the standard protocols (Stefan et al., 2000; Suppa et al., 2017; Wolters et al., 2003). Before protocol administration, the individual perceptual threshold for electrical median nerve stimulation was estimated, and electric stimulation during PAS was set at 300% of this value (see **2.6**). One hundred and eighty stimuli pairs were repeated with a frequency of 0.2 Hz. During PAS administrations, TMS was delivered at 110% rMT. The two PAS protocols differed only in the ISI between the two stimulations while keeping the other parameters constant (i.e., ISI of 25 ms for PAS_LTP_; ISI of 10 ms for PAS_LTD_). The choice of the parameters was made to find a good compromise between the duration of aftereffects, the duration of the protocol itself, and optimal parameters based on two published systematic reviews investigating the effects of PAS (Suppa et al., 2017; Wischnewski & Schutter, 2016). During PAS administration, participants were asked to count mentally the number of times the electric stimulation was delivered (i.e., 180), thus preventing sleepiness and keeping their attention high – a critical condition for the protocol’s effectiveness (Stefan et al., 2004).

To track the effects of PAS, MEPs and TEPs were acquired before (baseline, T0), immediately after (T1), and 30 minutes after PAS end (T2 – to investigate **H3**). In the TMS-EMG block, 30 trials were acquired. TMS-EEG blocks consisted of 120 trials each. Here, at T0 and T1, TMS was delivered at 110% rMT(suprathreshold) in one block and at 90% rMT (subthreshold) in the other (to investigate **H4**). At T2, we recorded only the block at suprathreshold intensity. The inter-pulse interval was randomly jittered between 3000 and 4000 ms in all the recording blocks acquired before and after PAS. During TMS-EMG blocks, TMS was delivered with the EEG cap on and under the same conditions and parameters of TMS-EEG recordings (i.e., noise masking was applied). See the **2.7** and **2.8** sections for further details. The TMS-EMG block lasted 3 minutes, while the TMS-EEG ones lasted 8 minutes each. During the TMS assessment, participants were at rest and instructed to keep their eyes open, looking at a fixation cross projected on the computer screen.

The order of the experimental sessions (i.e., PAS protocols) was counterbalanced across participants. TMS-EMG blocks were always delivered before TMS-EEG ones.

At the end of each session, three anatomical landmarks (nasion, left and right preauricular points) and the position of the 60 EEG channels were digitized for co-registration of the TMS-EEG data with the MRI template. On average, an experimental session lasted about 3 hours and 30 minutes.

### 2.6. TMS

Single-pulse TMS was delivered with an Eximia™ TMS stimulator (Nexstim™, Helsinki, Finland) using a biphasic focal figure-of-eight 70 mm coil. The stimulation target site was identified as the hotspot for the right APB muscle within the left M1. The location of the stimulation target was determined for each participant using a Navigated Brain Stimulation (NBS) system (Nexstim™, Helsinki, Finland) based on infrared-based frameless stereotaxy, allowing also accurate monitoring of the position and orientation of the coil and an online estimation of the distribution and intensity (V/m) of the intracranial electric field induced by the TMS. The coil was placed tangentially to the scalp and tilted 45° with respect to the midline (positioned perpendicular with respect to the stimulated cortical gyrus), inducing anterior-posterior (first phase)/posterior-anterior (second phase) currents within M1. Coil positioning was the same during EMG and EEG blocks.

TMS intensity was adjusted for each participant and session as a percentage of the rMT. rMT was preliminarily assessed in a short recording session before the experimental blocks using a parameter estimation by sequential testing (PEST) method (i.e., maximum-likelihood threshold-hunting procedure) (Awiszus, 2003; Dissanayaka et al., 2018). A sanity check ensured that 90% rMT stimulation did not induce corticospinal tract response: we assessed that no MEP was recorded in 10 consecutive trials from both APB and a cortical adjoining muscle (i.e., *first dorsal interosseus* – FDI) (Reijonen et al., 2020). If MEPs were present in one of these muscles at 90% rMT, motor hotspot searching was refined until the sanity check was fulfilled. Once the individual’s rMT value was determined, TMS intensity in the TMS-EEG blocks was set at 110% rMT or 90% rMT according to the experimental condition (see **Figure 1** and **H3**-**H4**). Considering the aim of TMS-EMG blocks (i.e., **H0)**, MEPs were recorded only at 110% rMT. Finally, during both PAS protocols, TMS was always administered at 110% rMT.

### 2.7. Electrical nerve stimulation

Median nerve stimulation during the PAS protocols employed a constant current stimulator (Digitimer DS7AH, Digitimer Ltd., Hertfordshire, UK). Surface electrodes were applied to stimulate the right-hand median nerve, exploiting a bipolar montage with the anode placed at the level of the wrist and the cathode proximal. The minimal intensity necessary to reliably elicit a sensation for each participant (based on self-report) was recognized as the perceptual threshold. Stimulation intensity during PAS was set at 300% of this value. The pulse width was set at 200 μS.

### 2.8. EEG recording

EEG data was continuously acquired from a 60-channel EEG cap (EasyCap, BrainProducts GmbH, Munich, Germany) using a sample-and-hold TMS-compatible system (Nexstim™, Helsinki, Finland). Two electrodes were placed over the forehead as ground and reference. Two additional electro-oculographic (EOG) channels were placed near the eyes (i.e., one above the right eyebrow and the other over the left cheekbone) to detect ocular artifacts due to eye movements and blinking (as done in: Bianco et al., 2023; Pisoni, Romero Lauro, et al., 2018; Romero Lauro et al., 2014). Noise masking was performed by continuously playing an audio track into earplugs created by shuffling TMS discharge noise to prevent the emergence of auditory evoked potentials (Russo et al., 2022). Noise masking volume was individually adjusted before each session to cover TMS clicks fully. Electrodes’ impedance was tested prior to each experimental session and kept below 5 kΩ. EEG signals were acquired with a sampling rate of 1450 Hz.

### 2.9. EMG recording

MEPs were recorded from the right-hand APB using Signal software (version 3.13) connected to a Digitmer D360 amplifier and a CED micro1401 A/D converter (Cambridge Electronic Devices, Cambridge, UK). Active electrodes (15 X 20 mm Ag-AgCl pre-gelled surface electrodes, Friendship Medical, Xi’an, China) were placed on the right thumb with a bipolar belly-tendon montage (i.e., active electrode over the muscle belly and reference electrode over the metacarpophalangeal joint of the thumb). The ground electrode was placed over the right head of the ulna. MEPs from the FDI muscle were recorded only during the sanity check for 90% rMT condition to assess the absence of corticospinal responses also in this second muscle (active electrode placed over the muscle belly and reference electrode over the metacarpophalangeal joint of the index). Before data acquisition, a visual check guaranteed that background noise did not exceed 20 μV. During TMS-EMG, participants also had noise masking to keep all recording conditions constant between EMG and EEG blocks. EMG signals were sampled (5000 Hz), amplified, band-pass filtered (10–1000 Hz) with a 50 Hz notch filter, and stored for offline analysis. Data was collected from 100 ms before to 200 ms after the TMS pulse (time window: 300 ms).

### 2.10. EEG preprocessing

EEG preprocessing was carried out in MATLAB (MathWorks, Natick, MA, USA) using EEGLAB (Delorme & Makeig, 2004) and TESA toolbox (Rogasch et al., 2017) functions. First, raw data was down-sampled to 725 Hz to reduce computational load. The continuous signal was re-referenced using an average reference, segmented in epochs starting 800 ms pre- and ending 800 ms post-TMS pulse, and baseline-corrected between −300 and −50 ms before TMS pulse. Single trials with excessive artifacts were rejected by visual inspection. The source-estimate-utilizing noise-discarding algorithm (SOUND, see Mutanen et al., 2018) implemented in TESA (Rogasch et al., 2017) was applied to attenuate extracranial noise coming from bad channels, exploiting a 3-layer spherical model with default parameters (λ = 0.1, as in Mutanen et al., 2018). Independent Component Analysis (FastICA, pop_tesa_fastica, ‘tanh’ contrast) was performed after Principal Component Analysis (PCA) compression to 30 components (pop_tesa_pcacompress). FastICA was solely applied to remove blinks and eye movements by visual inspection (Hernandez-Pavon et al., 2012). A semiautomatic signal space projection method for muscle artifact removal (SSP-SIR) was applied to suppress TMS-evoked muscle artifacts in the first 50 ms post-TMS (Mutanen et al., 2016). Epochs were band-pass filtered from 1 to 70 Hz and band-stop filtered from 48 to 52 Hz using a 4^th^-order Butterworth filter.

### 2.11. TEP extraction^1^

To narrow our investigation to the dynamics of left M1 local circuitry, we have computed the average of TEPs across a specified region of interest (ROI), including four electrodes under the stimulation coil or in correspondence with the scalp site of the cortical target, approximately C1, C3, C5, CP3, and FC3 (e.g., Costanzo et al., 2023; for a similar procedure see: Guidali et al., 2023; Lucarelli et al., 2025). First, the electrodes included in the ROI of each component were verified by visual inspection of the greatest positive (for P30 and P60) and negative (for N100) response amplitude in the time window selected for each TEP component from the baseline (i.e., T0) grand average of all participants, collapsing PAS_LTP_ and PAS_LTD_ sessions. Time windows of interest for P30, P60, and N100 components were selected according to the available literature on M1-TEP components elicited by both suprathreshold and subthreshold stimulations (e.g., Gordon et al., 2018; Lioumis et al., 2009; Lucarelli et al., 2025; Premoli, Castellanos, et al., 2014). They were: 20–35 ms for the P30, 55–70 ms for the P60, and 90-130 ms for the N100. Then, the clusters of electrodes were kept fixed among participants and, according to the ROI identified for each component, we extracted the individual amplitude value corresponding to the greatest positive (for P30 and P60) and negative (for N100) deflection in the aforementioned time intervals.

### 2.12. EMG preprocessing

Concerning EMG preprocessing, MEPs were analyzed offline using Signal software (version 3.13), following the standard preprocessing pipeline used in our laboratory (e.g., Guidali, Picardi, et al., 2023). At first, trials with artifacts (muscular or background noise) exceeding 200 µV in the 100 ms before the TMS pulse were automatically excluded. Then, MEP peak-to-peak amplitude was calculated in each trial between 5 ms and 60 ms from the TMS pulse. Trials in which MEP amplitude was smaller than 50 µV were excluded from the following analysis.

### 2.13. Planned statistical analysis

For our *positive control* condition (**H0**), MEP amplitude data was analyzed through planned comparisons using robust statistics (i.e., Yuen’s trimmed mean paired sample t-test, one-tailed, trimming level: 20%) (Mair & Wilcox, 2020; Yuen, 1974); in detail, according to our *a priori* hypothesis, we have tested that, for PAS_LTP_, MEP amplitude was higher after the administration of the protocol (T1) concerning the baseline (T0); for PAS_LTD_, we expected the reversed pattern (i.e., MEP amplitude lower than T0 after the PAS administration). For **H1**, PAS effects on TEP peak amplitude (i.e., P30 and P60) were separately analyzed through 2 X 2 within-subjects rmANOVA with factors ‘PAS protocol’ (PAS_LTP_, PAS_LTD_) and ‘Time’ (T0, T1).

For **H2**, PAS_LTD_ effects on N100 were assessed through robust statistics exploiting one-tailed Yuen’s trimmed mean paired sample t-test (Mair & Wilcox, 2020; Yuen, 1974), comparing N100 amplitude before (T0) and after (T1) the administration of PAS_LTD_.

For **H3**, the temporal profile of PAS aftereffects on MEP, P30, P60, and N100 amplitude was investigated through 3 X 2 within-subjects rmANOVA with factors ‘PAS protocol’ (PAS_LTP_, PAS_LTD_) and ‘Time’ (T0, T1, T2).

Finally, for **H4**, possible effects of supra- or subthreshold intensity on P30 and P60 amplitude in the PAS_LTP_ were investigated. Given the rationale of our *a priori* hypothesis (see **1**), this analysis would have been conducted if **H1** had shown significant modulation of P60 amplitude after PAS_LTP_ administration. Here, for each component, we would calculate the ratio of T1 peak amplitude over T0 (*PAS_LTP_ effect*). Then, the ‘post-pre amplitude’ ratio was used as the dependent variable in a 2 X 2 within-subjects rmANOVA with factors ‘Intensity’ (90%, 110%) and ‘Component’ (P30, P60).

In all our rmANOVAs, significant main effects and interactions were further explored with post-hoc tests by applying Tukey’s correction for multiple comparisons. If data sphericity was not confirmed by Mauchly’s test, the Greenhouse–Geisser correction has been applied. Partial eta-squared (*η_p_^2^* – for rmANOVAs) and Cohen’s *d* (for t-tests) were reported as effect size values. The mean ± standard error (SE) was reported for each variable. Statistical significance was set at *p* < .02. The normality of our data distributions was tested using the Shapiro-Wilks test and Q-Q plot assessment. If normality is not achieved, to make the distribution closer to normality, we have transformed the raw data with three commonly used transformations for continuous variables: (*a*) square root [i.e., 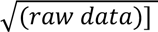, (*b*) base-ten logarithmic [i.e., log_10_(raw data)], and (*c*) inverse transformation [i.e., 1/(raw data)]. To account for possible negative values, as well as values between 0 and 1, when applying these transformations, we added a constant to the raw data values, thus anchoring the minimum of our distribution(s) to 1 (Osborne, 2010). Then, we have selected among these three transformations the one showing the best fit to a normal distribution (i.e., the transformed distribution presents values of an excess kurtosis between −2 and 2 and skewness between −1 and 1; the distribution which values fell into these ranges, being closer to 0, was selected –George & Mallery, 2019). Statistical analyses were performed using the Jamovi software (The Jamovi Project, 2025), R Studio (R Core Team, 2020), and Fieldtrip (Oostenveld et al., 2011).

## 3. RESULTS

### 3.1. Final sample and TMS-EEG preprocessing

Forty-five healthy participants took part in the study (25 females, mean age ± SD: 23.6 ± 3.9 years; mean education ± SD: 14.7 ± 2.1 years; mean Edinburgh score ± SD: 84.7 ± 17.2%). Considering the preplanned exclusion criteria, 15 participants were not included in the analysis due to the following reasons: (*i*) 5 participants did not complete the experiment due to technical or personal issues, (*ii*) 2 participants were excluded because MEP or TEP amplitude exceeds 3 SD from the mean of the group, and (*iii*) 8 participants presented TMS-EEG cleaned data with SNR < 1.5 in at least one experimental condition.

Hence, the analyzed sample comprised 30 participants (19 females, mean age ± SD: 24 ± 4.2 years; mean education ± SD: 15.1 ± 2.2 years; mean Edinburgh score ± SD: 84 ± 18.4%). Detailed information on mean participants’ rMT, TMS intensities, perceptual thresholds for PAS electric stimulation, number of ICA and SSP-SIR components removed during EEG preprocessing are reported in **Supplemental Tables S1** and **S2**. TEP grand averages and P30, P60, and N100 topographies in the different experimental conditions are reported in **Figure 2**. Considering the grand average of all participants (see components’ topographies depicted in **Figure 2**), the following ROIs were considered for peak extraction: Cz, C2, CP2, and CP4 for the P30; CP1, CP3, CP5, and P3 for the P60 and the N100. Even if there is no consensus in literature on the precise cluster of electrodes to be selected for each M1-TEP components, the topographies of our components are consistent with previous TMS-EEG literature, suggesting that we have correctly recorded the neural components of interest evoked by M1 stimulation (e.g., Biabani et al., 2019; e.g., Lucarelli et al., 2025; Premoli, Castellanos, et al., 2014; Zazio et al., 2021). The mean activity in the selected ROIs is reported in **Supplemental Figure S1.**

**Figure 2.**
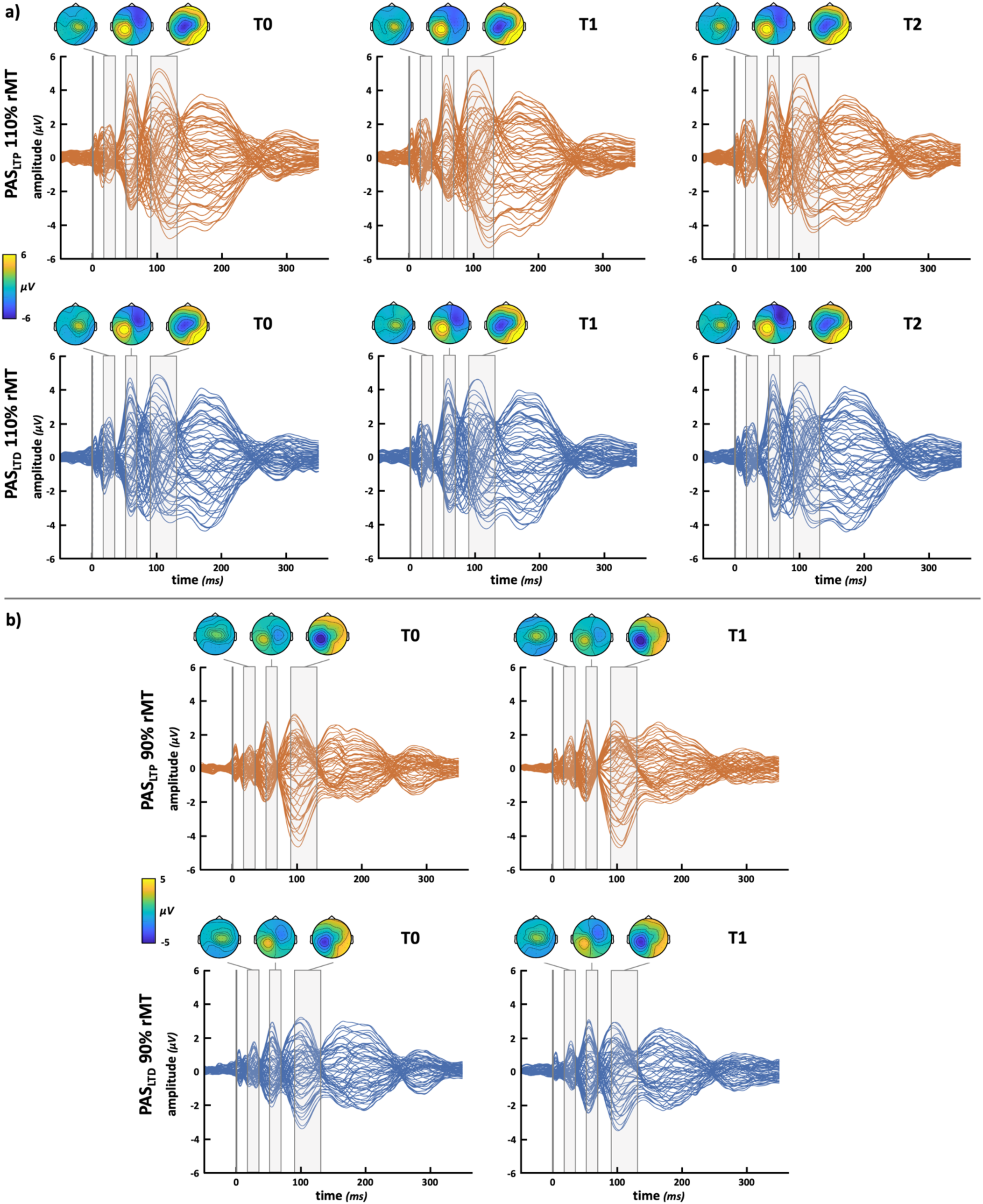
TEP grand averages. Butterfly plots of TEP grand average recorded before (T0), immediately after (T1), and 30 minutes (T2) from PAS administration (orange traces = PAS_LTP_; blue traces = PAS_LTD_). **(a)** suprathreshold (i.e., 110% rMT) conditions. **(b)** subthreshold (i.e., 90% rMT) conditions. Grey-shaded areas over the plots show time windows of P30, P60, and N100 components extraction and their topographies. The voltage scale used for topographies changes between 110% (panel **a**) and 90% rMT (panel **b**) conditions.

MEP and P30 amplitudes did not follow a normal distribution. Transforming them with the base-ten logarithm made their distribution closer to normality and within the preplanned ranges. Hence, analyses on MEP and P30 were conducted on log-transformed raw values.

This work received Stage 1 in-principle acceptance (IPA) on 15/01/2024. The IPA version of the manuscript is publicly available on Open Science Framework (OSF) Registries (https://osf.io/detjc). Raw data, datasets, analyses, and scripts can be found on OSF (https://osf.io/48fh3/).

### 3.2. Registered Analyses

#### 3.2.1. MEP amplitude (H0 – *positive control*)

We found that (log-transformed) MEP amplitudes were significantly higher after the administration of the PAS_LTP_ (2.75 ± 0.06) than in baseline (2.58 ± 0.04; *t*_17_ = 3.61, *p* = .001, *d* = .7). Conversely, MEPs amplitude was significantly reduced after the LTD-inducing protocols (2.57 ± 0.04, *vs*. T0: 2.67 ± 0.04; *t*_17_ = −2.84, *p* = .006, *d* = .41; **Figure 3**; **Supplemental Figure S2a** for the single-subject ratio of T1 MEP amplitude over T0 – i.e., *PAS effect*). This pattern follows the one expected from previous literature, confirming that PAS_LTP_ and PAS_LTD_ protocols can successfully induce LTP and LTD phenomena detectable at the corticospinal level after administration.

**Figure 3.**
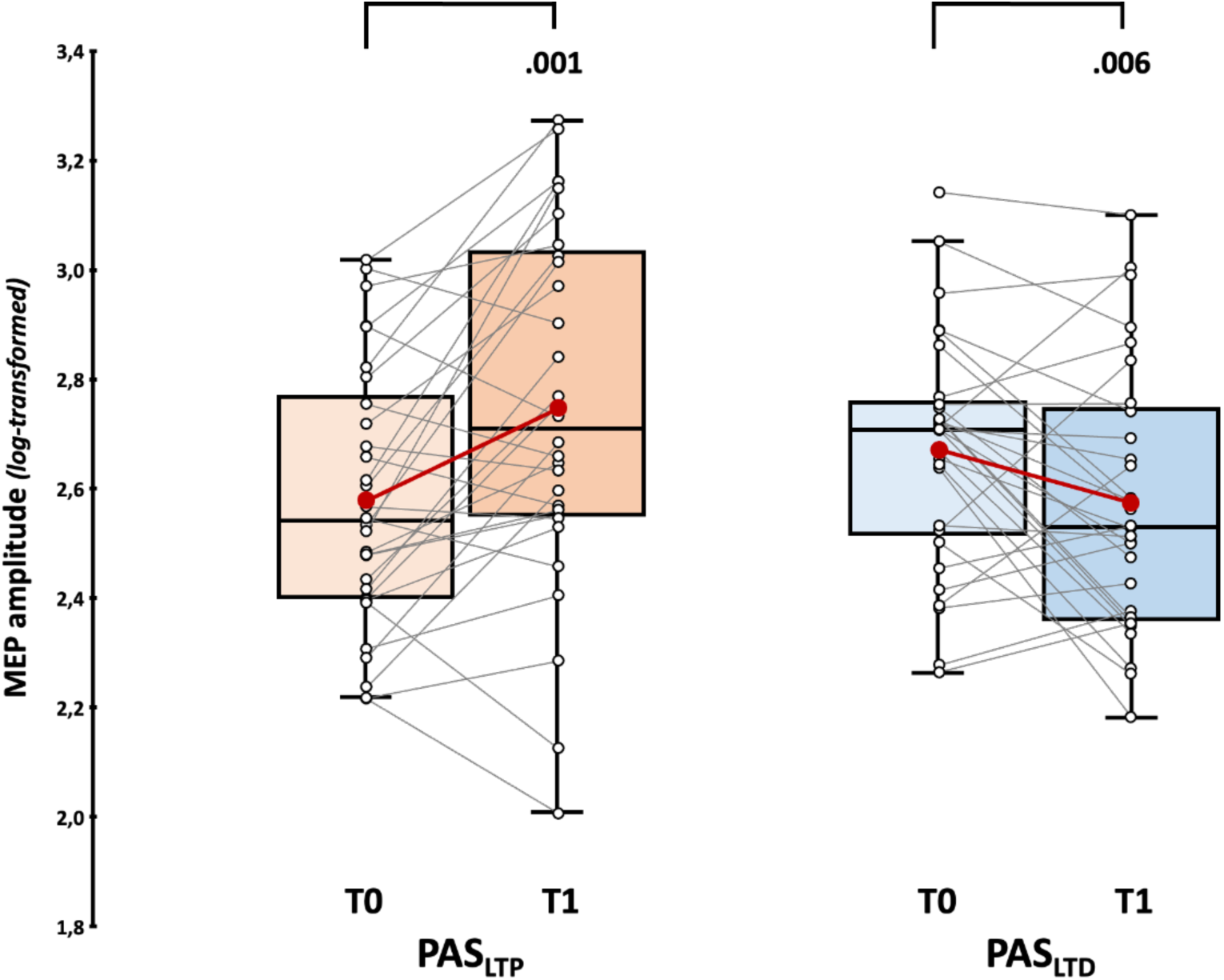
Corticospinal results. Upper panel: (log-transformed) MEP amplitude assessed before (T0) and immediately after (T1) PAS_LTP_ (orange boxplots) and PAS_LTD_ (blue boxplots) administration. Red dots and lines indicate the means of the distributions. The center line denotes their median values. Black-and-white dots and grey lines show individual participants’ scores. The box contains the 25^th^ to 75^th^ percentiles of the dataset. Whiskers extend to the largest observation falling within the 1.5 * inter-quartile range from the first/third quartile. Significant *p* values of Yuen’s trimmed means paired sample t-tests are reported.

#### 3.2.2. PAS effects on P30 and P60 components (H1)

Considering the rmANOVA on (log-transformed) P30 amplitude, we found a significant effect of factor ‘Time’ (*F*_1,29_ = 6.23, *p* = .018, *η_p_^2^* = .16) and, crucially, of the ‘PAS protocol’ X ‘Time’ interaction (*F*_1,29_ = 11.84, *p* = .002, *η_p_^2^* = .29). Post-hoc showed that P30 amplitude was significantly higher after PAS_LTP_ administration (0.58 ± 0.05) compared to the baseline (0.47 ± 0.04; *t*_29_ = 3.22, *p_tukey_* = .016, *d* = .59). Notably, no differences occurred between T0 (0.52 ± 0.04) and T1 (0.5 ± 0.04) for PAS_LTD_ (*t*_29_ = 1.62, *p_tukey_* = .384, *d* = .3; **Figure 4a, Supplemental Figure S2b** for the single-subject ratio of T1 P30 amplitude over T0 – i.e., *PAS effect*). The main effect of ‘PAS protocol’ did not reach statistical significance (*F*_1,29_ = 0.2, *p* = .661, *η_p_^2^* < .01).

**Figure 4.**
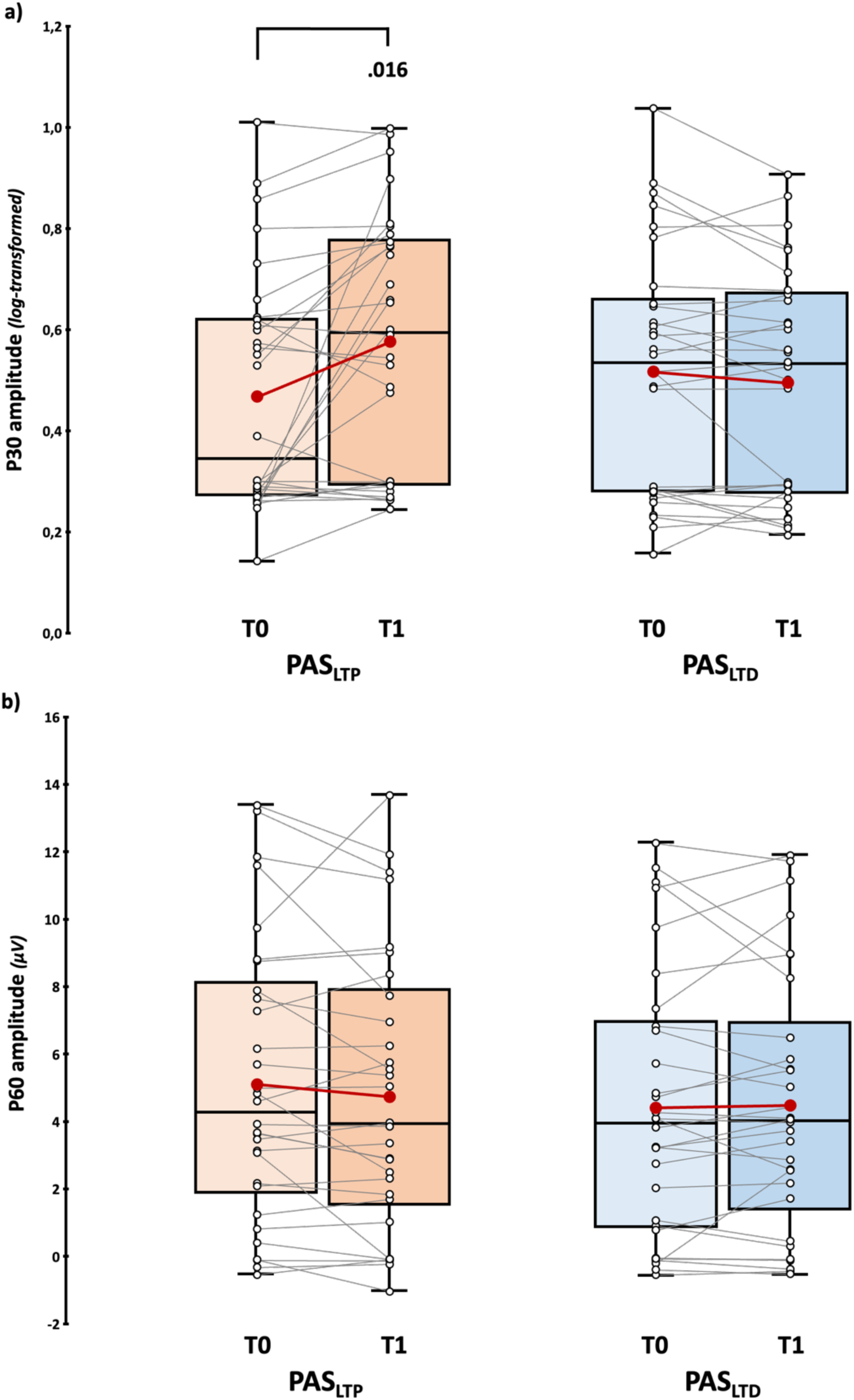
P30 and P60 results. (log-transformed) P30 (**a**) and P60 amplitude (**b**) assessed before (T0) and immediately after (T1) PAS_LTP_ (orange boxplots) and PAS_LTD_ (blue boxplots) administration. Red dots and lines indicate the means of the distributions. The center line denotes their median values. Black-and-white dots and grey lines show individual participants’ scores. The box contains the 25^th^ to 75^th^ percentiles of the dataset. Whiskers extend to the largest observation falling within the 1.5 * inter-quartile range from the first/third quartile. Significant *p* values of Tukey corrected post-hoc comparisons are reported.

Considering the P60, neither the main effects (‘PAS protocol’: *F*_1,29_ = 1.72, *p* = .199, *η_p_^2^* = .06; ‘Time’: *F*_1,29_ = 0.84, *p* = .367, *η_p_^2^* = .03) nor their interaction (*F*_1,29_ = 1.42, *p* = .243, *η_p_^2^* = .05) reached statistical significance (**Figure 4b, Supplemental Figure S2c** for the single-subject ratio of T1 P60 amplitude over T0 – i.e., *PAS effect*).

Hence, considering immediate PAS effects on early components, we found that only the P30 was selectively modulated after PAS_LTP_. Importantly, given the absence of significant effects on P60 amplitude, the possible influence of reafferent processing on TEPs components (i.e., **H4**) was exploratorily conducted and reported in the **3.3.1** section.

#### 3.2.3. PAS effects on N100 (H2)

Considering the N100, robust paired sample t-test showed that its amplitude was significantly higher after PAS_LTD_ administration (−5.49 ± 0.76 μV) compared to baseline (−4.73 ± 0.66 μV; *t*_17_ = 2.31, *p* = .017, *d* = .49). For PAS_LTP_, we found no pre-post difference (T0: −5.38 ± 0.75 μV, *vs.* T1: −5.59 ± 0.87 μV; *t*_17_ = 0.21, *p* = .419, *d* < .01; **Figure 5, Supplemental Figure S2d** for the single-subject ratio of T1 N100 amplitude over T0 – i.e., *PAS effect*).

**Figure 5.**
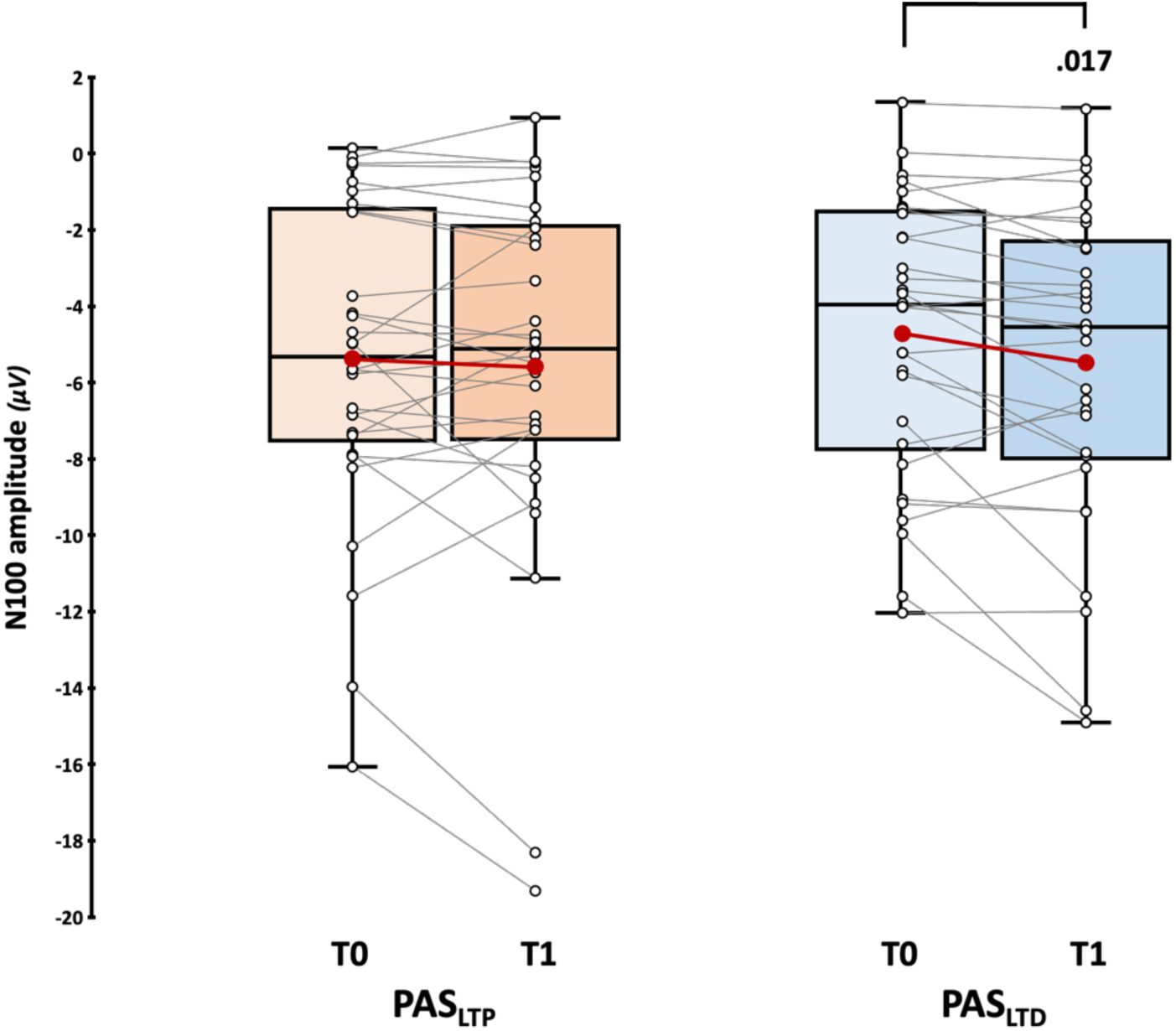
N100 results. Upper panel: N100 amplitude assessed before (T0) and immediately after (T1) PAS_LTP_ (orange boxplots) and PAS_LTD_ (blue boxplots) administration. Red dots and lines indicate the means of the distributions. The center line denotes their median values. Black-and-white dots and grey lines show individual participants’ scores. The box contains the 25^th^ to 75^th^ percentiles of the dataset. Whiskers extend to the largest observation falling within the 1.5 * inter-quartile range from the first/third quartile. Significant *p* values of Yuen’s trimmed means paired sample t-tests are reported.

#### 3.2.4. Temporal evolution of PAS effects (H3)

Considering the temporal evolution of corticospinal effects, the rmANOVA showed only a significant ‘PAS protocol’ X ‘Time’ interaction (*F*_2,58_ = 12.83, *p* < .001, *η_p_^2^* = .31), confirming that T1 MEPs presented opposite patterns after PAS_LTP_ and PAS_LTD_ administration. Nevertheless, post-hoc comparisons did not show any statistically significant effects when T2 values (PAS_LTP_: 2.68 ± 0.05, PAS_LTD_: 2.58 ± 0.05) were compared to T0 or T1 ones (all *t*s < 2.68, all *p*s > .11), suggesting that PAS modulations on corticospinal excitability assessed after 30 minutes from the end of the PAS showed greater variability than immediate effects (**Figure 6a**, all T2 *vs.* T1 and T0 post-hoc comparisons are reported in **Supplemental Table S3**). Main effects did not reach significance (‘PAS protocol’: *F*_1,29_ = 5.1, *p* = .032, *η_p_^2^* = .15; ‘Time’: *F*_2,58_ = 1.05, *p* = .356, *η_p_^2^* = .04).

**Figure 6.**
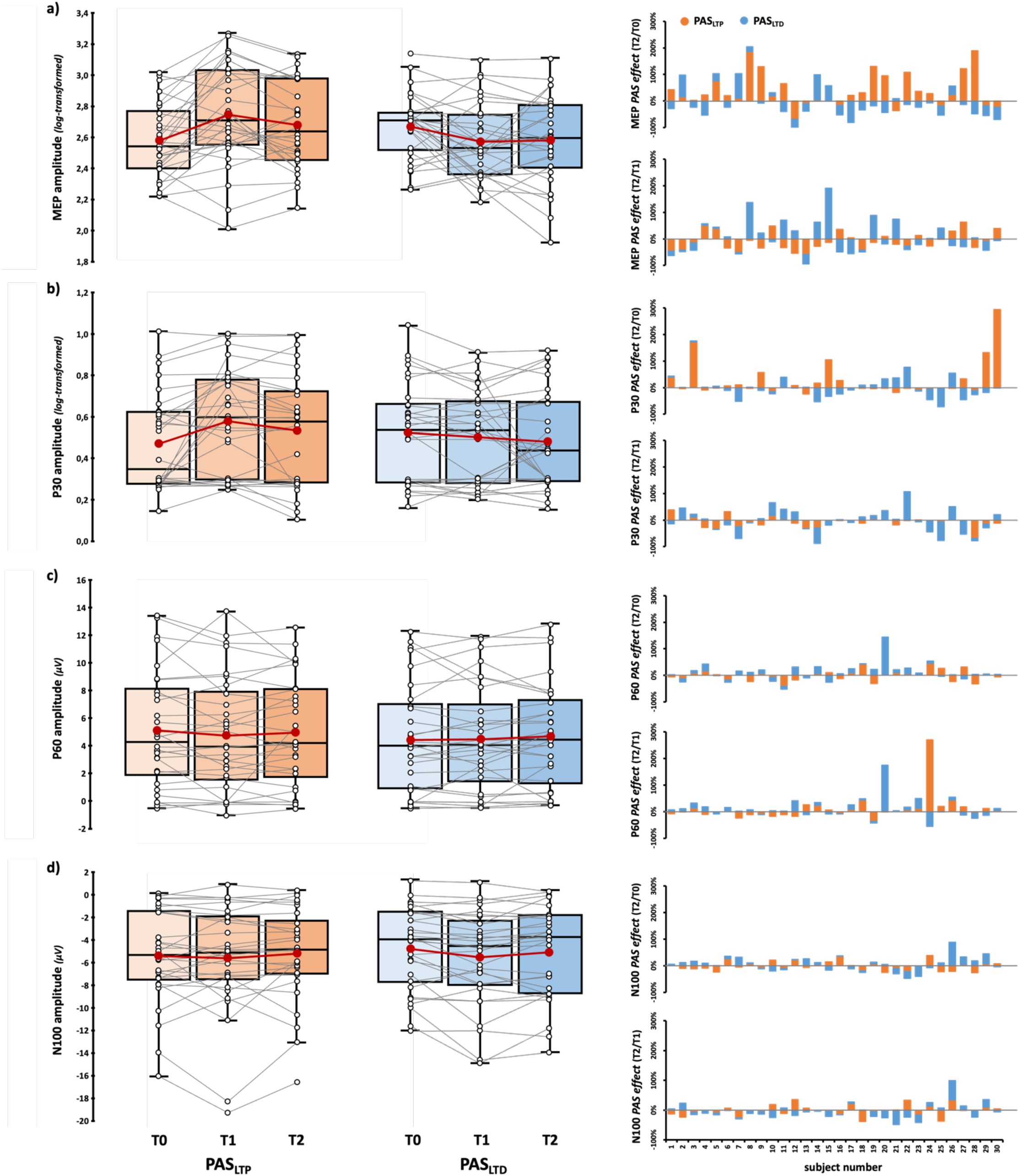
Temporal patterns of PAS effects. Left panels: (log-transformed) MEP (**a**), (log-transformed) P30 (**b**), P60 (**c**), and N100 (**d**) amplitude assessed before (T0) immediately after (T1), and 30 minutes after (T2) PAS_LTP_ (orange boxplots) and PAS_LTD_ (blue boxplots) administration. Red dots and lines indicate the means of the distributions. The center line denotes their median values. Black-and-white dots and grey lines show individual participants’ scores. The box contains the 25^th^ to 75^th^ percentiles of the dataset. Whiskers extend to the largest observation falling within the 1.5 * inter-quartile range from the first/third quartile. Right panels: *PAS effects* at different timing (i.e., the ratio between T2 and T0 and between T2 and T1) at the single-subject level according to the two protocols (orange bars: PAS_LTP_, blue bars: PAS_LTD_).

A similar pattern was also found for the P30, where only the ‘PAS protocol’ X ‘Time’ interaction was significant (*F*_2,58_ = 6.13, *p* = .004, *η_p_^2^* = .18; ‘PAS protocol’: *F*_1,29_ = 0.94, *p* = .341, *η_p_^2^* = .03; ‘Time’: *F*_2,58_ = 2.78, *p* = .07, *η_p_^2^* = .09), highlighting P30 enhancement after PAS_LTP_ administration. Post-hoc comparisons did not show differences when T2 (log-transformed) amplitudes (PAS_LTP_: 0.53 ± 0.05, PAS_LTD_: 0.48 ± 0.04) were compared to T0 or T1 ones (all *t*s < 2.12, all *p*s > .304; **Figure 6b, Supplemental Table S3**).

For the P60, neither main effects (‘PAS protocol’: *F*_1,29_ = 1.13, *p* = .297, *η_p_^2^* = .04; ‘Time’: *F*_2,58_ = 0.96, *p* = .39, *ηp^2^* = .03) nor their interaction reached statistical significance (*F*_2,58_ = 0.89, *p* = .415, *η_p_^2^* = .03, **Figure 6c**). For the N100, only the main effect of factor ‘Time’ reached statistical significance (*F*_2,58_ = 4.46, *p* = .016, *η_p_^2^* = .13). Post-hoc shows only a tendency towards significance for the T2 *vs.* T1 comparison (*t*_29_ = 2.58, *p_tukey_* = .04, *d* = .37) suggesting that, regardless of the specific protocol, N100 amplitudes at T2 (−5.11 ± 0.72 μV) were less negative than values obtained at T1 (−5.54 ± 0.77 μV, **Figure 6d**). ‘PAS protocol’ (*F*_1,29_ = 0.59, *p* = .45, *η_p_^2^* = .02) and ‘PAS protocol’ X ‘Time’ interaction (*F*_2,58_ = 1.08, *p* = .345, *η_p_^2^* = .04) were not statistically significant.

### 3.3. Exploratory Analyses

#### 3.3.1. M1-TEP peaks at subthreshold intensity and effects of MEP reafferent processing (H4)

Given that no modulation was found on the P60, **H4** and analyses on TEPs recorded at 90% rMT were carried out exploratively. Nine subjects considered for the planned analyses have at least one block of recording at 90% where SNR was < 1.5, and they were excluded from this set of analyses. Thus, we ran analyses on 90% rMT conditions on a sub-sample of 21 participants (13 females, mean age ± SD: 23.4 ± 2.8 years; mean education ± SD: 14.8 ± 2.2 years; mean Edinburgh score ± SD: 83.9 ± 20.6%), making **H4** likely under- powered (see **Table 1, Sampling plan** column) and the results obtained should be interpreted with caution. **Figure 2b** depicts TEP grand averages in the 90% rMT conditions.

First, we explored whether TEP components (i.e., P30, P60, and N100) recorded at 90% rMT differed after PAS administration, conducting a series of ‘PAS protocol’ X ‘Time’ rmANOVAs, as the ones performed for testing **H1**. For all TEP components, analyses were conducted on log-transformed values. None of these analyses showed statistically significant effects or interactions, suggesting that M1-TEP peaks were not modulated by the two PAS protocols when recorded at a subthreshold intensity (all *F*s < 1.5, all *p*s > .235, **Figure 7a**, **Supplemental Table S4**). Then, we run a rmANOVA on the ratio of T1 peak amplitude over T0 recorded during PAS_LTP_ (*PAS_LTP_ effect*) to explore the influence of reafferent processing on TEP components (**H4)**.

**Figure 7.**
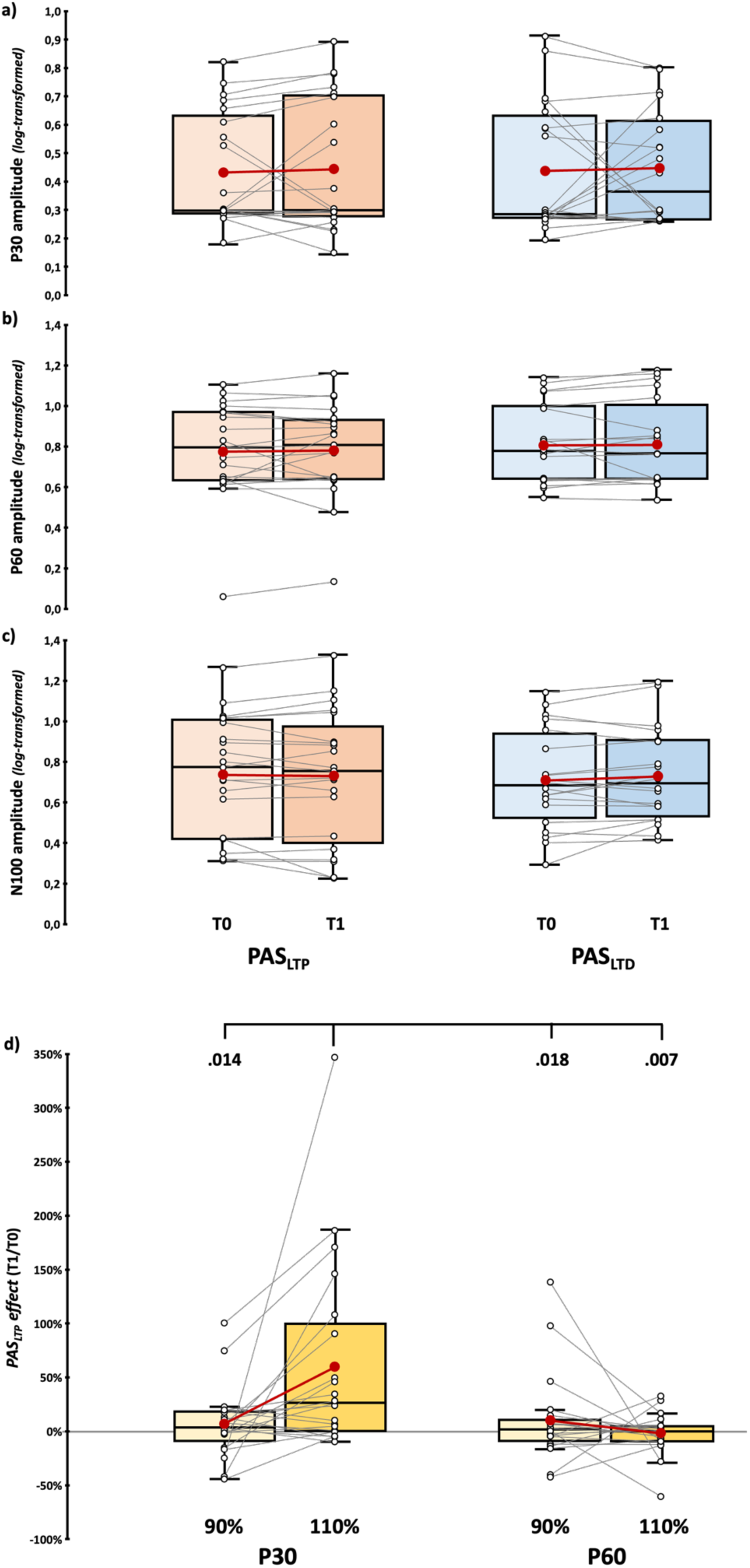
Results of subthreshold analyses on TEPs. (**a, b, c**) (log-transformed) P30, P60, and N100 amplitude assessed before (T0) and immediately after (T1) PAS_LTP_ (orange boxplots) and PAS_LTD_ (blue boxplots) administration in subthreshold blocks. **(d)** *PAS_LTP_ effect* (i.e., the ratio between T1 and T0 amplitude) for P30 (left panel) and P60 components (right panel) recorded sub- and supra-threshold. Significant *p* values of planned comparison t-tests are reported. In the box-and-whiskers plots, red dots and lines indicate the means of the distributions. The center line denotes their median values. Black-and-white dots and grey lines show individual participants’ scores. The box contains the 25^th^ to 75^th^ percentiles of the dataset. Whiskers extend to the largest observation falling within the 1.5 * inter-quartile range from the first/third quartile.

This analysis showed a significant ‘Intensity’ X ‘Component’ interaction (*F*_1,20_ = 7.81, *p* = .011, *η_p_^2^* = .28) and a main effect of factor ‘Component’ (*F*_1,20_ = 7.57, *p* = .012, *η_p_^2^* = .27). Planned t-tests showed that P30 *PAS_LTP_ effect* at 110% rMT (60.1 ± 19.5%) was significantly higher than at 90% rMT (7.2 ± 7.2%; *t*_20_ = 2.69, *p* = .014, *d* = .59) and P60 *PAS effects* at 110% rMT (−1.7 ± 4.2%; *t*_20_ = 3.1, *p* = .006, *d* = .66) and 90% rMT (10.3 ± 9%; *t*_20_ = 2.59, *p* = .018, *d* = .57; **Figure 7b**). This pattern suggests that the P30 recorded suprathreshold was the only TEP component modulated by PAS_LTP_.

#### 3.3.2. Correlations between MEP and TEP modulations

To further explore possible associations between corticospinal and cortical modulations, we run a series of Spearman’s correlations between the ratio of T1 MEP amplitude over T0 (MEP *PAS effect*) and the ratio of T1 TEP peaks amplitude over T0 (i.e., P30, P60, and N100 *PAS effect*), separated for PAS_LTP_ and PAS_LTD._ No significant correlations were found (PAS_LTP_, all *ρ*s < .16, all *p*s > .397, **Figure 8a**; PAS_LTD_, all *ρ*s < .03, all *p*s > .866, **Figure 8b**).

**Figure 8.**
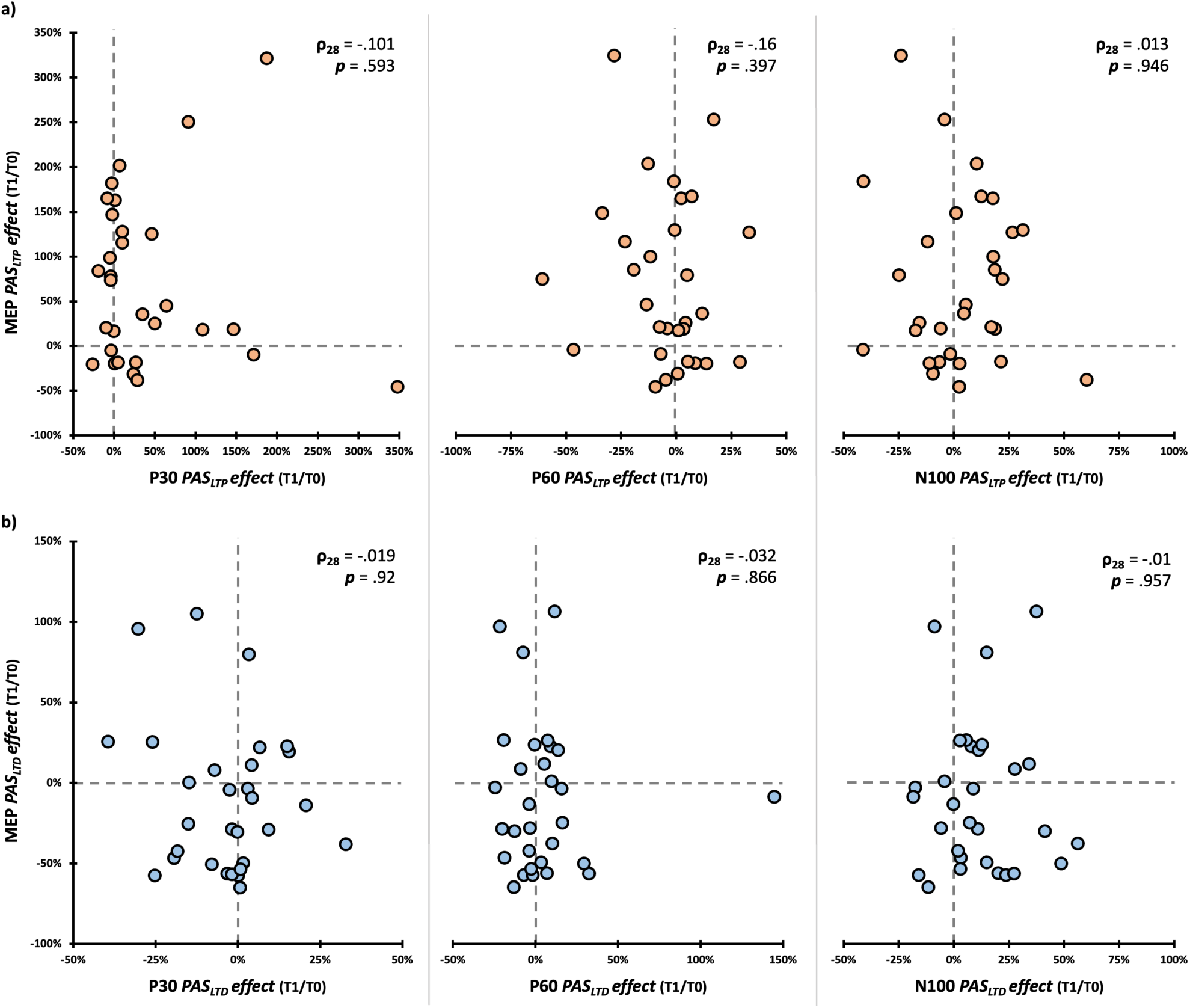
Correlation between PAS-induced corticospinal and cortical modulations. Scatterplot between MEP *PAS effect* (i.e., the ratio between T1 and T0 amplitude) obtained for the excitatory (**a**) and inhibitory (**b**) protocols and P30, P60, and N100 *PAS effects* found after the same PAS condition. In the upper corners, we reported Spearman correlation coefficients and related p-values.

#### 3.3.3. Responders/non-responders’ ratio and predictability of PAS modulations from baseline assessment

As the literature highlights (e.g., Fratello et al., 2006; Minkova et al., 2019), PAS protocols suffer from a potentially consistent share of non-responders. **Figure 9a** reported percentages of PAS responders and non-responders at T1 at the group and single-subject level according to the four variables modulated after PAS administration (i.e., MEP and P30 amplitude after PAS_LTP_, MEP and N100 amplitude after PAS_LTD_; see **3.2.1**, **3.2.2**, **3.2.3**). To account for near-zero values, we adopted a conservative approach (Leodori et al., 2021), considering as ‘responders’ participants who presented *PAS effect* values (i.e., the ratio of T1 amplitude over T0) greater than 5% for MEP_LTP_, P30_LTP_, and N100_LTD,_ or smaller than −5% for MEP_LTD_. At the corticospinal level, PAS_LTP_ responders were 70% (21/30) of our sample, while 57% (17/30) were PAS_LTD_ responders. Considering cortical variables (i.e., P30 and N100), 57% (17/30) were PAS_LTP_ responders, while 60% (18/30) were PAS_LTD_ ones (**Figure 9a**).

**Figure 9.**
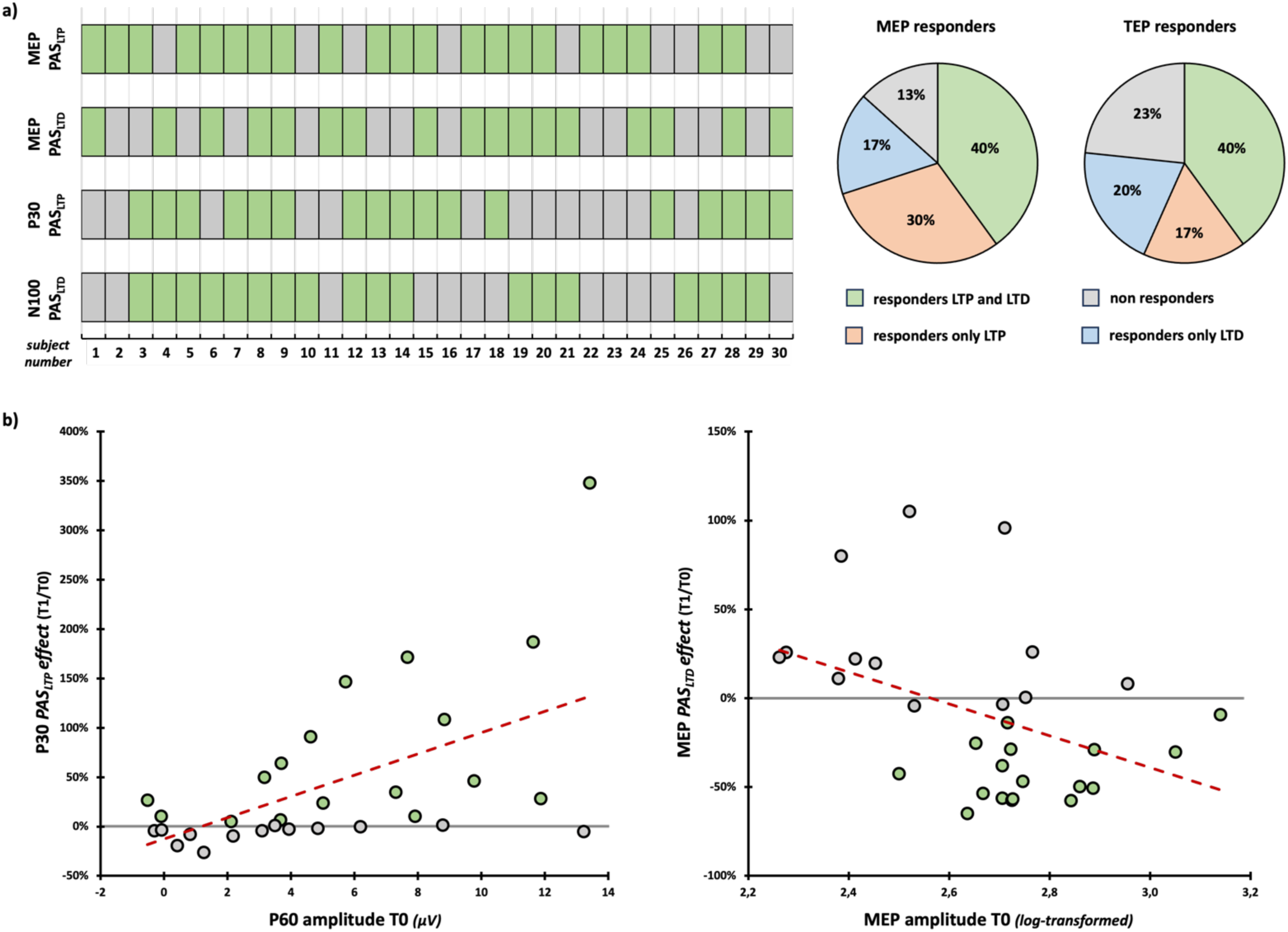
**(a)** *PAS responder and non-responder characterization.* Single-subject distribution of responders and non-responders according to the four variables modulated after the two PAS protocols. **(b)** Scatterplots between *PAS effects* (i.e., the ratio between T1 and T0 amplitude) and (log-transformed) MEP and P60 amplitude. Red dashed lines indicate the linear regression’s fitted line. Grey dots indicate non-responders and green dots indicate PAS responders.

To better explore PAS responders’ and non-responders’ profiles, we investigated whether neurophysiological variables (i.e., rMT, MEP, P30, P60, and N100 amplitude) recorded in baseline (T0) could predict MEP_LTP_, MEP_LTD_, P30_LTP_, and N100_LTD_ *PAS effects*. To this aim, we run a series of generalized linear regression models with the aforementioned variables (distribution: Gaussian; link function: Identity; Gallucci, 2019). Results showed that P60 baseline values predicted the magnitude of P30 modulations after PAS_LTP_ administration (model fit: R^2^ = 0.32, χ^2^_1_ = 12.99, *p* < .001): namely, the greater the P60 at T0, the greater the P30 amplitude gain after PAS_LTP_ (B = 0.11, SE = .03, z = 3.6., *p* < .001). Similar to P60, MEP amplitude at T0 significantly predicted PAS_LTD_ modulations at the corticospinal level (model fit: R^2^ = 0.17, χ^2^_1_ = 1.05, *p* = .015): i.e., the greater the MEP amplitude before PAS_LTD_ administration, the greater the magnitude of MEP inhibition (B = −0.9, SE = 0.37, z = −2.43, *p* = .015; **Figure 9b**). None of the other generalized linear regression models showed statistically significant effects (all χ^2^s < 3.6, all *p*s > .032; **Supplemental Table S5**).

## 4. DISCUSSION

In the present Registered Report study, we aimed to investigate the cortical correlates of PAS-induced LTP and LTD, leveraging TMS-EEG and assessing possible modulations of TEP components recorded from M1 after PAS administration. To date, PAS effects have been evaluated predominantly through corticospinal measures (i.e., MEPs – Suppa et al., 2017). However, since MEPs are a mixed measure of central and peripheral signal conduction, they only indirectly measure PAS-induced changes at the cortical level. Therefore, TMS-EEG recordings are essential to capture associative plasticity effects occurring at the cortical level to better ground PAS neurophysiological mechanisms, optimizing its effectiveness and its use in clinical settings (e.g., Baroni et al., 2024; Suppa et al., 2017; Wischnewski & Schutter, 2016).

Overall, our results showed that, beyond confirming MEPs enhancement and inhibition after PAS_LTP_ and PAS_LTD_, the excitatory protocol induced a significant modulation of the P30 component, and at the same time, the inhibitory one selectively modulated the N100, suggesting that these two TEP components could reflect cortical biomarkers of PAS-induced associative plasticity.

### 4.1. PAS effects at the corticospinal level

The results of our *positive control* analysis on PAS-induced modulations at the corticospinal level confirmed the effectiveness of the two protocols: PAS_LTP_ significantly increased MEP amplitude, while PAS_LTD_ led to a reduction, in line with previous literature on PAS-induced Hebbian associative plasticity (Carson & Kennedy, 2013; Suppa et al., 2017; Wischnewski & Schutter, 2016). However, both PAS protocols did not always produce MEP increasing or decreasing immediately after their administration: only 12 out of the 30 participants responded to both PAS_LTP_ and PAS_LTD_ protocols, while 3 participants were classified as ‘fully’ non-responders since they did not show any excitability changes following both PAS protocols. On the one hand, this pattern confirms previous findings of high interindividual variability in PAS effects on corticospinal excitability (e.g., Fratello et al., 2006; Lahr et al., 2016; López-Alonso et al., 2014; Minkova et al., 2019), reinforcing the idea that plasticity induction tracked peripherally may depend on intrinsic neurophysiological differences at the cortical level that need further characterization. On the other hand, this result suggests carefulness when the term ‘spike-timing dependent plasticity’ is used to describe PAS modulations and corticospinal aftereffects. Indeed, by taking advantage of the within-subject design of our study, we show that less than half of our sample (40%) responded simultaneously to both protocols, as instead expected if PAS-induced modulations followed classic STDP rules observed in vitro or animal models (e.g., Brzosko et al., 2019; Caporale & Dan, 2008; Dan & Poo, 2004). This evidence is further corroborated at the cortical level, where we found a similar percentage of responders and non-responders (**Figure 9a**).

### 4.2. PAS effects on positive M1-TEP components (P30 and P60)

Concerning cortical measures, we successfully replicated the significant increase in P30 amplitude following PAS_LTP_ previously reported by Costanzo and coworkers (2023). P30 has been associated with local excitatory neurotransmission (Mäki & Ilmoniemi, 2010), and the evidence that PAS_LTD_ does not modulate its amplitude in the opposite direction suggests that the excitatory mechanisms reflected by the component are insensitive to plastic changes induced by the LTD protocol and are somewhat specific to LTP phenomena, supporting the role of segregated mechanisms underlying these effects. Our results are also consistent with previous studies using TMS-EEG to assess plastic changes in cortical excitability, where increases in the early TEP components were observed following excitatory neuromodulation interventions (Cruciani et al., 2023; Esser et al., 2006; Pisoni, Mattavelli, et al., 2018). Hence, our findings confirm that P30 may reflect a reliable marker of early cortical excitability enhancement within the motor network after PAS_LTP_ administration.

Conversely, P60 amplitude remained unchanged following PAS_LTP_ and PAS_LTD._ This result diverges from Costanzo et al. (2023) and prior research reporting similar modulations for the two components following TMS protocols known to modulate M1 excitability (Cash et al., 2017). Some studies previously highlighted the potential role of reafferent signals from activated muscles in the presence of MEPs (i.e., when using supra-threshold intensities – Fecchio et al., 2017; Komssi et al., 2004; Petrichella et al., 2017). For instance, in a study by Petrichella and colleagues (2017), trials eliciting MEPs presented higher P60 amplitude than trials where MEPs were not evoked, suggesting that somatosensory reafference may influence modulations reported for this component (Petrichella et al., 2017). Albeit exploratory, our comparative analysis with data recorded at a subthreshold intensity (i.e., 90% rMT) seems to rule out a bias driven by reafference in the observations on P60 amplitude before *vs.* after the two PAS protocols, given that the T1-T0 ratio recorded in supra- and subthreshold conditions is not differently modulated. In contrast to previously reported findings (Costanzo et al., 2023), we argue that P60 may not be a reliable marker of PAS-induced plasticity.

Nevertheless, our exploratory analyses indicate that P60 might be a sensitive predictor for PAS_LTP_ responsiveness at the cortical level. P60 amplitude reflects glutamatergic transmission (Belardinelli et al., 2021), and it has been reported that glutamatergic agonists facilitate LTP-like effects following neuromodulation, lowering the threshold for LTP induction (Cohen & Abraham, 1996). This might explain why, in our study, individual baseline P60 amplitude levels significantly predicted enhancement of the P30 component linked to early excitatory cortical activity. P60 amplitude is attenuated by short-latency afferent inhibition (SAI), a phenomenon involving cortico-cortical signal transmission between the primary somatosensory cortex (S1) and M1 (Ferreri et al., 2012). SAI is inversely correlated with LTP-like PAS-induced effects and explains a fair share of inter-individual variability in PAS outcomes (Guerra et al., 2020; Murase et al., 2015). Collectively, these findings warrant future investigations to consider individual differences in baseline P60 amplitude when interpreting PAS_LTP_ outcomes.

Regarding analyses on TEPs recorded at 90% rMT, these subthreshold assessments failed to identify modulations on P30 amplitude similar to the ones found using suprathreshold intensities. This result is interesting, considering that the P30 is reliably recorded in our 90% rMT blocks (**Figure 2b** and **7a**) and should not be influenced by MEP reafference. However, considering the neurophysiology of PAS-induced plasticity (e.g., Lamy et al., 2010; Meunier et al., 2007; Stefan et al., 2002; Suppa et al., 2017) it has to be noted that PAS_LTP_ is thought to potentiate M1 interneuron synapses, like S1-M1 connections (Carson & Kennedy, 2013; Hamada et al., 2014; Ni et al., 2019; Stefan et al., 2002), as well as M1 pyramidal neurons, which can alter their discharge properties. At the same time, we know that, at variance with subthreshold stimulation, suprathreshold pulses over M1 recruits also nearby and interconnected regions, like the postcentral gyrus and somatosensory cortices (Bestmann et al., 2004; Shitara et al., 2011; Siebner et al., 2022) and that, by definition, subthreshold TMS over M1 activates pyramidal neurons located in deeper cortical layers and projecting to the corticospinal tract to a lesser extent than suprathreshold stimulation (McColgan et al., 2020; Siebner et al., 2022). Importantly, TEPs reflect a compound signal from different cortical sources (Hernandez-Pavon, Veniero, et al., 2023). Hence, we propose that suprathreshold TMS could better capture PAS-induced modulations, as reflected by P30 patterns, by activating a greater neuronal population both within M1 and in its proximal connections. In this regard, we stress that the P30 reflects excitatory activity involving multiple cortical sources related to M1 local circuitry (Farzan & Bortoletto, 2022; Mäki & Ilmoniemi, 2010), and we argue that the suprathreshold stimulation (110% of the rMT) may better activate pyramidal neurons within M1 than the subthreshold stimulation (90% of the rMT), making it more suitable for highlighting the plastic modulation induced by PAS_LTP_ at the cortical level. In the future, it will be interesting to assess the effects of both PAS protocols on TEP components peaking earlier than P30, like immediate TEPs (i-TEPs), which were proposed to be the most genuine cortical evoked response by TMS, directly reflecting the synchronized excitation of pyramidal neurons in the targeted M1 (Beck et al., 2024).

### 4.3. N100 as a cortical marker of PAS_LTD_

Concerning PAS_LTD_, in line with our hypothesis, we reported a significant increase of N100 amplitudes following this protocol, potentially reflecting the upregulation of GABAergic activity, which is thought to contribute to LTD-like neuromodulatory effects (Cruciani et al., 2023). Indeed, this result expands previous TMS-EEG reports on PAS effects, as the modulations reported in the seminal work of Huber and colleagues (2008) were not in a time window compatible with the N100 component, while Costanzo and coworkers (2023) only investigated the effects of the LTP-inducing protocol. In line with the literature (e.g., Bonnard et al., 2009; Casula et al., 2014; Premoli et al., 2018; Premoli, Rivolta, et al., 2014; Rogasch et al., 2013), our results reinforce the specificity of N100 as an inhibitory marker, specifically suitable for tracking PAS_LTD_ effects.

### 4.4. Temporal dynamics of PAS aftereffects

Concerning the temporal dynamics of PAS_LTP_ effects, we obtained a complex pattern of results across both MEPs and P30 amplitude modulations. Specifically, T2 measures – compared to baseline and immediate post-PAS effects – suggest that changes in corticospinal (i.e., MEPs) and cortical (i.e., P30) facilitation markers may persist after PAS, yet with much more variability across subjects (**Figure 6a, right panels** and non-significant T1 vs. T2 comparisons – **Supplemental Table S2**). Although our results do not demonstrate that the effects of PAS_LTP_ persist 30 minutes after administration, they are nonetheless indicative that, at least in a cohort of participants, PAS_LTP_ aftereffects might last beyond the duration of the protocol, potentially due to consolidation mechanisms.

Considering PAS_LTD_, we did not observe any long-lasting effects when analyzing the temporal evolution of cortical and corticospinal measures after its administration. Moreover, this inhibitory protocol seems more prone to interindividual variability immediately after administration (Wischnewski & Schutter, 2016). Taken together, such findings suggest that the excitatory effects of PAS_LTP_ are stronger than the depressing effects of PAS_LTD_. The reasons behind this difference are not fully understood. We can speculate that LTD effective induction may rely more on the pre-existing state of M1 excitability at baseline (Goldsworthy et al., 2014) or may be contrasted by homeostatic mechanisms counteracting activity suppression (Abraham, 2008). Along with this idea, we found that MEP amplitude at T0 significantly predicts PAS_LTD_ modulations at the corticospinal level. This evidence supports that the M1 functional state may critically determine the variability of PAS_LTD_ corticospinal effects.

### 4.5. Methodological considerations and future directions

Although we showed that PAS induces plasticity at both the cortical and corticospinal levels (as reflected in modulations of MEPs and TEP components, respectively), cortical and corticospinal excitability effects appear to be independent (i.e., no correlations were observed between them at baseline and after the PAS; **Figure 8**). The independence of these measures supports the notion that M1-TEP and MEP patterns reflect distinct functional states of motor cortex activation (e.g., Biabani et al., 2021; Guidali et al., 2025; Guidali, Zazio, et al., 2023; Madsen et al., 2019). However, it must be noted that, in the present work, MEP and TEP recordings occurred in separate blocks due to technical reasons related to our setup. Hence, future works integrating EEG and EMG in the same recording block could better link PAS-induced cortical and corticospinal modulations. Another point that deserves some reflection is that in the 90% rMT recordings, we could not obtain an SNR > 1.5 for the whole sample, so analyses involving these data could be underpowered. Although stimulating with an intensity below the rMT is a valid methodological approach for testing hypotheses that require disambiguating cortical effects from corticospinal contribution (e.g., reafferent signals), this approach impacts the reliability of the protocol since a subthreshold stimulation cannot always be able to elicit a cortical response large enough to distinguish it from baseline. Hence, stimulation not only does not activate the corticospinal tract but also does not elicit a stable response of the neural populations below the coil. This technical consideration should inform future research that wants to test the cortical effects of PAS through TMS-EEG and, to a greater extent, TMS-EEG studies adopting subthreshold stimulation conditions to investigate cortical effects of neuromodulatory protocols.

Finally, effect sizes found across our analyses suggest that PAS_LTP_ aftereffects are more robust than PAS_LTD_ ones and, hence, that the LTP protocol is more effective and likely replicable, at least adopting the parameters of the present work. The evidence that LTP can be more easily inducted than LTD with PAS is confirmed by evidence from modified versions targeting other sensory and cognitive systems, as well as cortico-cortical pathways. These protocols commonly found modulations reassembling LTP induction rather than LTD, suggesting that the former direction of Hebbian plasticity is more easily inducible than the latter (for reviews, see: Di Luzio et al., 2024; Guidali et al., 2021b; Hernandez-Pavon, San Agustín, et al., 2023). Further, it must be noted that an important factor potentially hindering a reliable induction of PAS effects could be the adoption of fixed parameters, especially the timing between electrical stimulation and TMS over M1 (i.e., 25 ms for PAS_LTP_ and 10 ms for PAS_LTD_). It has been shown that adjusting the ISI between paired stimulations according to the individual N20 latency, i.e., the first cortical component of the median nerve somatosensory-evoked potential, could represent an effective method for enhancing the reliability of the neuromodulation outcomes at the cortico-spinal level (e.g., Müller-Dahlhaus et al., 2008; Suppa et al., 2017; Ziemann et al., 2004). Future studies should deepen whether individualized parameters can improve the reliability of PAS effects also at the cortical level.

## 5. CONCLUSION

In conclusion, taking advantage of the Registered Report format, we demonstrate that P30 and N100 M1-TEP components act as markers of LTP and LTD induction after PAS administration. Our findings could inform future studies investigating associative plasticity with TMS-EEG, as well as clinical investigations taking advantage of PAS protocols targeting the motor system. At the same time, they also highlight the need for further research to deepen factors leading to the high inter-individual variability of cortical and corticospinal PAS aftereffects and participants’ different sensibility to LTP or LTD induction.

## ACKNOWLEDGMENTS

We thank Gloria Galbiati, Danila Ippoliti, Sara Meazzi, Elisa Morini, Laura Sassi, and Margherita Varenna for their valuable help in data collection.

## DATA AVAILABILITY STATEMENT

Registered Report snapshot, Stage 1/2 versions of the manuscript, raw data, datasets, analyses, and scripts can be found on Opens Science Framework - OSF (https://osf.io/48fh3/)

## CRedIT AUTHOR CONTRIBUTION

**Eleonora Arrigoni:** conceptualization, methodology, investigation, software, visualization, writing – original draft

**Nadia Bolognini:** methodology, supervision, writing – review & editing

**Alberto Pisoni:** methodology, supervision, validation, resources, funding acquisition, writing – review & editing

**Giacomo Guidali:** conceptualization, methodology, validation, data curation, investigation, software, visualization, writing – original draft

## Footnotes

1 The following section was rephrased during *Stage 2* to enhance the transparency and reproducibility of our peak extraction procedure

