## Supplemental Materials for "Cortical markers of PAS-induced long-term potentiation and depression in the motor system:A TMS-EEG Registered Report"

### – SUPPLEMENTARY MATERIALS –

| Protocol | rMT | 110% condition and PAS | 90% condition | sensory threshold (electrical stimulation) | PAS electrical stimulation intensity |
|---|---|---|---|---|---|
| **$PAS_{LTP}$** | 43.9 ± 8.5% | 48.4 ± 9.3% | 39.1 ± 7.6% | 3.16 ± 0.83 mA | 9.49 ± 2.37 mA |
| **$PAS_{LTD}$** | 43.3 ± 8.6% | 47.6 ± 9.5% | 38.3 ± 7.5% | 3.14 ± 0.66 mA | 9.09 ± 2.26 mA |

**Supplemental Table S1.** TMS and electrical stimulator's mean ± standard deviation – SD of parameters used in the experimental sessions.

| | | $PAS_{LTP}$ | | | | | $PAS_{LTD}$ | | | | |
|---|---|---|---|---|---|---|---|---|---|---|---|
| | | 110% | | | 90% | | 110% | | | 90% | |
| | | T0 | T1 | T2 | T0 | T1 | T0 | T1 | T2 | T0 | T1 |
| **ICA components removed** | ***Mean*** | 2,07 | 2,10 | 2,17 | 2,20 | 2,03 | 2,07 | 2,23 | 2,07 | 2,13 | 2,20 |
| | ***SD*** | 1,08 | 0,76 | 0,75 | 1,13 | 0,67 | 1,05 | 0,94 | 0,98 | 0,82 | 0,81 |
| **SSP-SIR components removed** | ***Mean*** | 2,90 | 2,79 | 2,86 | 1,97 | 1,90 | 2,83 | 2,79 | 2,72 | 2,17 | 2,24 |
| | ***SD*** | 0,88 | 0,76 | 0,86 | 1,02 | 1,03 | 0,76 | 0,76 | 0,64 | 1,00 | 1,11 |

**Supplemental Table S2.** Mean ± SD number of ICA and SSP-SIR components removed during EEG preprocessing in the different experimental sessions.

| Variable | Protocol | Comparison | Statistics |
|---|---|---|---|
| **(log-transformed) MEP** | $PAS_{LTP}$ | T2 *vs.* T0 | $t_{29}$ = 2.68, $p_{tukey}$ = .11 |
| | | T2 *vs.* T1 | $t_{29}$ = -2.24, $p_{tukey}$ = .25 |
| | $PAS_{LTD}$ | T2 *vs.* T0 | $t_{29}$ = -2.25, $p_{tukey}$ = .245 |
| | | T2 *vs.* T1 | $t_{29}$ = 0.3, $p_{tukey}$ = .99 |
| **(log-transformed) P30** | $PAS_{LTP}$ | T2 *vs.* T0 | $t_{29}$ = 2.12, $p_{tukey}$ = .304 |
| | | T2 *vs.* T1 | $t_{29}$ = -2.14, $p_{tukey}$ = .297 |

| | | | |
|---|---|---|---|
| | $PAS_{LTD}$ | T2 *vs.* T0 | $t_{29}$ = -1.47, $p_{tukey}$ = .687 |
| | | T2 *vs.* T1 | $t_{29}$ = -0.67, $p_{tukey}$ = .984 |

**Supplemental Table S3.** T2 *vs.* T0 and T1 post-hoc comparisons for statistically significant 'PAS protocol' X 'Time' interactions found for **H3** analyses.

| **Variable** | **Factor** | **Statistics** |
|---|---|---|
| **(log-transformed) P30** | PAS protocol | $F_{1,19}$ = 0.03, $p$ = .867, $\eta_p^2$ < .01 |
| | Time | $F_{1,19}$ = 0.5, $p$ = .489, $\eta_p^2$ = .03 |
| | PAS protocol X Time | $F_{1,19}$ = 0.07, $p$ = .799, $\eta_p^2$ < .01 |
| **(log-transformed) P60** | PAS protocol | $F_{1,19}$ = 0.75, $p$ = .396, $\eta_p^2$ = .04 |
| | Time | $F_{1,19}$ = 0.03, $p$ = .864, $\eta_p^2$ < .01 |
| | PAS protocol X Time | $F_{1,19}$ < 0.01, $p$ = .99, $\eta_p^2$ < .01 |
| **(log-transformed) N100** | PAS protocol | $F_{1,19}$ = 0.18, $p$ = .674, $\eta_p^2$ = .01 |
| | Time | $F_{1,19}$ = 0.22, $p$ = .647, $\eta_p^2$ = .01 |
| | PAS protocol X Time | $F_{1,19}$ = 1.5, $p$ = .235, $\eta_p^2$ = .07 |

**Supplemental Table S4.** Results from the rmANOVAs conducted on M1-TEP peaks extracted in the 90% rMT conditions.

| **Dependent variable** | **Predictor** | **Generalized linear regression model fit** |
|---|---|---|
| **MEP *$PAS_{LTP}$ effect*** | rMT | $R^2$ = 0.01, $\chi^2_1$ = 0.21, $p$ = .628 |
| | (log-transformed) MEP | $R^2$ < 0.01, $\chi^2_1$ = 0.16, $p$ = .675 |
| | (log-transformed) P30 | $R^2$ < 0.01, $\chi^2_1$ = 0.1, $p$ = .74 |
| | P60 | $R^2$ = 0.01, $\chi^2_1$ = 0.21, $p$ = .629 |
| | N100 | $R^2$ = 0.14, $\chi^2_1$ = 3.6, $p$ = .031 |
| **MEP *$PAS_{LTD}$ effect*** | rMT | $R^2$ = 0.01, $\chi^2_1$ = 0.04, $p$ = .666 |
| | **(log-transformed) MEP** | **$R^2$ = 0.17, $\chi^2_1$ = 1.05, $p$ = .015** |
| | (log-transformed) P30 | $R^2$ = 0.02, $\chi^2_1$ = 0.11, $p$ = .463 |
| | P60 | $R^2$ = 0.03, $\chi^2_1$ = 0.19, $p$ = .339 |
| | N100 | $R^2$ < 0.01, $\chi^2_1$ < 0.01, $p$ = .955 |
| **P30 *$PAS_{LTP}$ effect*** | rMT | $R^2$ < 0.01, $\chi^2_1$ = 0.02, $p$ = .864 |
| | (log-transformed) MEP | $R^2$ = 0.01, $\chi^2_1$ = 0.22, $p$ = .568 |
| | (log-transformed) P30 | $R^2$ = 0.11, $\chi^2_1$ = 2.02, $p$ = .068 |
| | **P60** | **$R^2$ = 0.32, $\chi^2_1$ = 12.99, $p$ < .001** |

| | | |
|---|---|---|
| | N100 | $R^2 = 0.04$, $\chi^2_1 = 0.69$, $p = .306$ |
| **N100 *PAS$_{LTD}$ effect*** | rMT | $R^2 = 0.01$, $\chi^2_1 = 0.01$, $p = .623$ |
| | (log-transformed) MEP | $R^2 < 0.01$, $\chi^2_1 < 0.01$, $p = .956$ |
| | (log-transformed) P30 | $R^2 = 0.11$, $\chi^2_1 = 0.11$, $p = .069$ |
| | P60 | $R^2 < 0.01$, $\chi^2_1 < 0.01$, $p = .813$ |
| | N100 | $R^2 = 0.01$, $\chi^2_1 = 0.01$, $p = .578$ |

**Supplemental Table S5.** Results from the generalized linear regression models (function: Gaussian, link: Identity) ran to explore the predictability of T0 variables for PAS aftereffects (i.e., ratio of T1 values over T0). Significant ($p < .02$) effects are reported in bold.

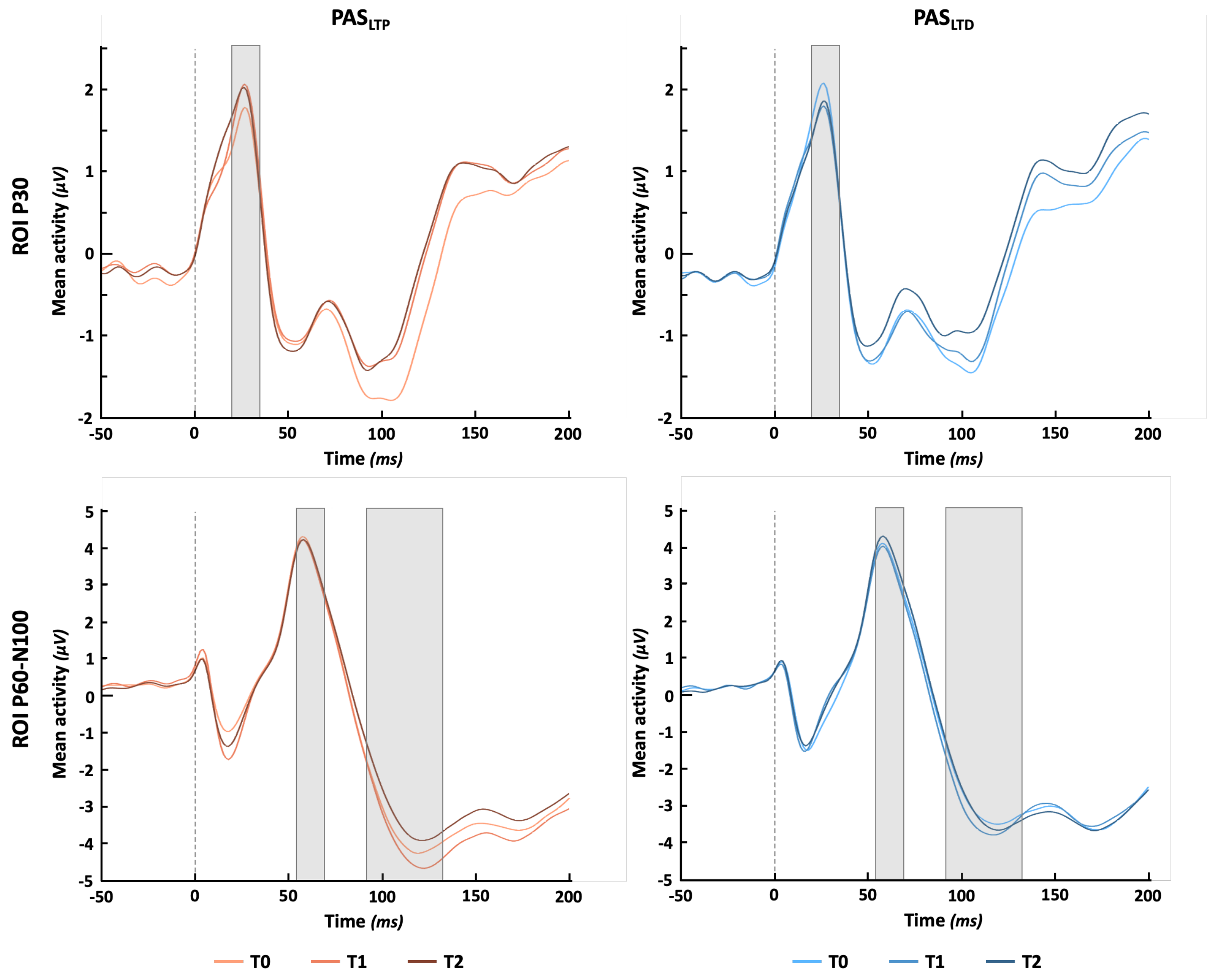


**Supplemental Figure S1.** Mean activity in the two ROIs selected for peak extraction (upper panels: P30 ROI: Cz, C2, CP2, CP4 electrodes, lower panels: P60 and N100 ROI: CP1, CP3, CP5, P3 electrodes) in 110% rMT conditions. Grey-shaded areas over the plots show time windows of P30, P60, and N100 components extraction.

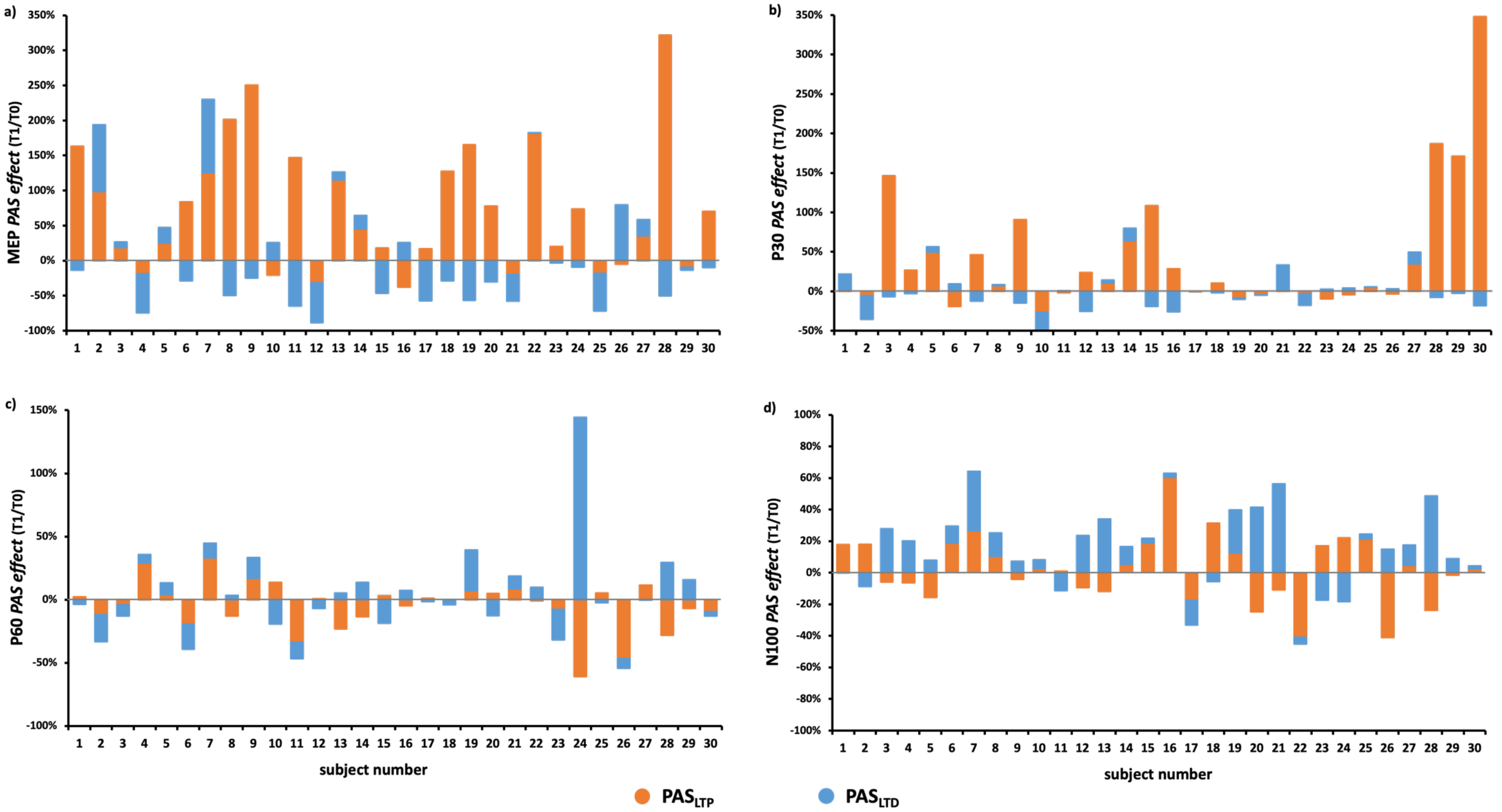


**Supplemental Figure S2.** *PAS effect* (i.e., the ratio between T1 and T0 amplitude) for MEPs (**a**), P30 (**b**), P60 (**c**), and N100 (**d**) components at the single-subject level according to the two protocols (orange bars: $PAS_{LTP}$, blue bars: $PAS_{LTD}$).
